# Transposable element expression at unique loci in single cells with CELLO-seq

**DOI:** 10.1101/2020.10.02.322073

**Authors:** Rebecca V Berrens, Andrian Yang, Christopher E Laumer, Aaron TL Lun, Florian Bieberich, Cheuk-Ting Law, Guocheng Lan, Maria Imaz, Daniel Gaffney, John C Marioni

## Abstract

The role of Transposable Elements (TEs) in regulating diverse biological processes, from early development to cancer, is becoming increasing appreciated. However, unlike other biological processes, next generation single-cell sequencing technologies are ill-suited for assaying TE expression: in particular, their highly repetitive nature means that short cDNA reads cannot be unambiguously mapped to a specific locus. Consequently, it is extremely challenging to understand the mechanisms by which TE expression is regulated and how they might themselves regulate other protein coding genes. To resolve this, we introduce CELLO-seq, a novel method and computational framework for performing long-read RNA sequencing at single cell resolution. CELLO-seq allows for full-length RNA sequencing and enables measurement of allelic, isoform and TE expression at unique loci. We use CELLO-seq to assess the widespread expression of TEs in 2-cell mouse blastomeres as well as human induced pluripotent stem cells (hiPSCs). Across both species, old and young TEs showed evidence of locus-specific expression, with simulations demonstrating that only a small number of very young elements in the mouse could not be mapped back to with high confidence. Exploring the relationship between the expression of individual elements and putative regulators revealed surprising heterogeneity, with TEs within a class showing different patterns of correlation, suggesting distinct regulatory mechanisms.

## Introduction

Gene expression signatures characterize distinct cellular states, and changes in these signatures are associated with both normal development and disease. Consequently, understanding the complete picture of expression within a single cell is important for understanding these processes ^1^. Short read single-cell RNA-sequencing (scRNA-seq) has recently emerged as a vital tool for characterising the transcriptomes of individual cells, facilitating projects such as the Human Cell Atlas ^2,3^. However, the majority of scRNA-seq protocols use short-read sequencing and, moreover, are biased towards tagging either the 3’ or 5’ end of a transcript. Consequently, the ability to profile individual isoforms, to assay allele specific expression and, importantly, to study the expression of individual transposable elements in single-cells has generally remained challenging ^4–9^.

The expression of TEs is being increasingly recognised as playing a key role in multiple biological processes, with a recent focus being their function in early mammalian development ^10^. More generally, half of the mammalian genome is estimated to consist of repetitive DNA elements, with TEs contributing to 45% of the human and 37.5% of the mouse genome ^11,12^. TEs can be classified based upon their method of transposition, with Class 1 elements (including long and short interspersed elements (LINEs and SINEs) and Long Terminal Repeats (LTRs)) that transpose via RNA intermediates and Class 2 elements, or DNA transposons, that transpose via a DNA intermediate through a so-called cut-and-paste mechanism ^3^. Most TEs are silenced, existing as fragmented copies in the genome. However, 1-2% of TEs are young, full-length and active in both the mouse (e.g. SINE-B1, L1MdA, L1MdF, MERVL and IAPs elements) and the human (e.g., L1HS, SVA-E, AluYb8 and HERV-int) genome ^13–17^. Epigenetic modifications repress TEs in most somatic cells but they become transcriptionally active during early development and global epigenetic reprogramming ^18,19^. Recent work on LINE expression during early development has hinted at a role for retrotransposons in regulating gene expression via an unknown mechanism ^20,21^. However, due to the highly repetitive nature of TEs across the genome, short cDNA reads cannot be unambiguously mapped to a specific TE locus-of-origin. Therefore, measures of TE expression have been aggregated across multiple elements at a family or subfamily level, limiting the ability to understand the mechanisms by which they might regulate the expression of messenger RNAs and hence normal development.

To address this, we have developed a novel experimental protocol, for single CELl LOng read RNA sequencing (CELLO-seq) (Figure 1A), and an associated computational framework, which combines the benefits of existing scRNA-seq protocols with advances in long-read sequencing as implemented by Oxford Nanopore Technologies (ONT) and Pacific Biosciences (PacBio). Our approach uses long Unique Molecular Identifiers (UMIs) to counteract the high error rate associated with ONT reads relative to Illumina sequencing. By profiling mouse blastomeres from 2-cell stage embryos and a larger set of human Induced Pluripotent Stem Cells (hiPSCs) CELLO-Seq uncovers unexpected heterogeneity in expression of individual elements within specific repeat classes. Moreover, these differences are associated with different upstream epigenetic regulators, suggesting different modes of regulation.

**Figure 1.**
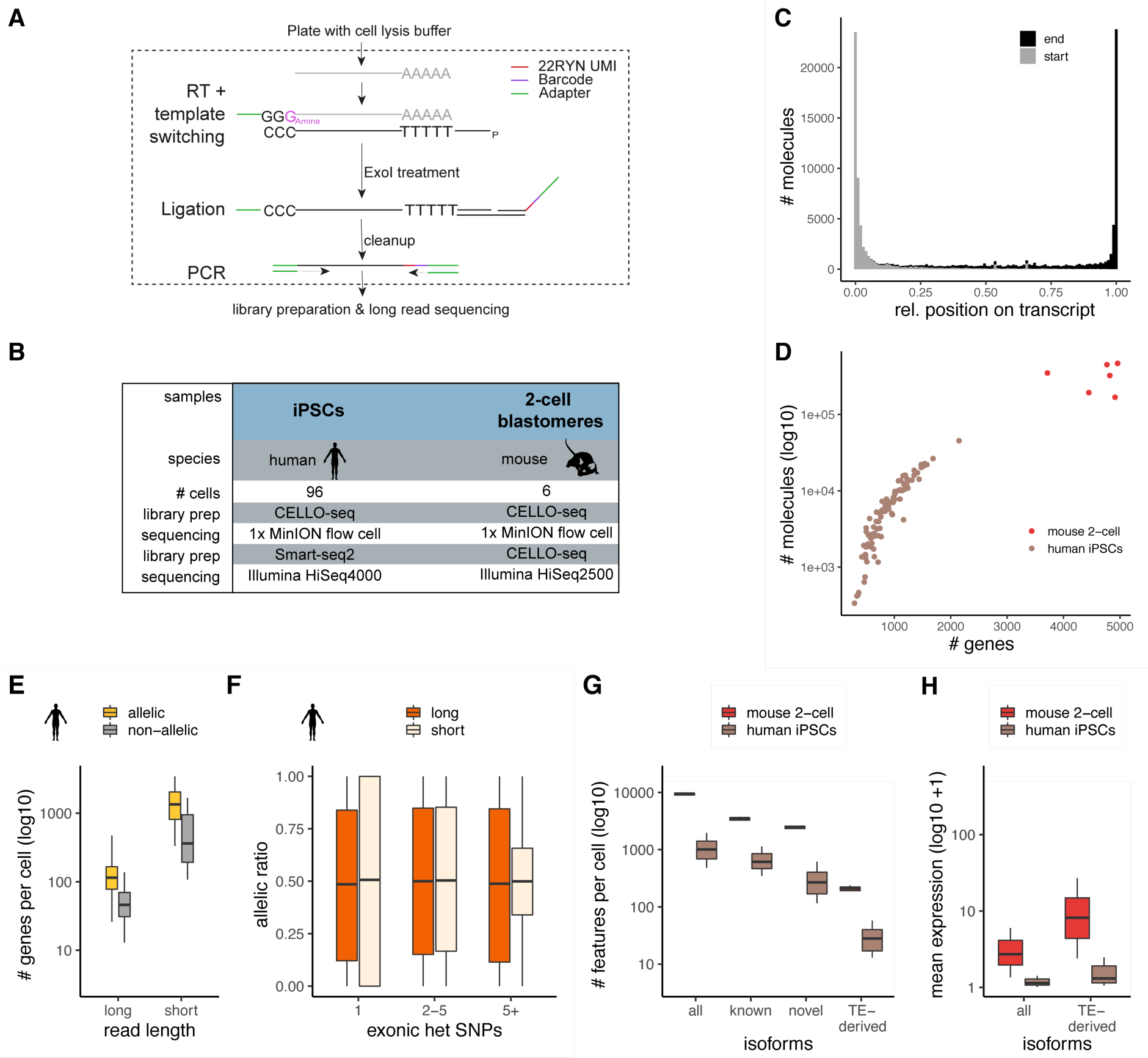
CELLO-seq overview and ability to study allelic and isoform expression. (A) CELLO-seq protocol overview. Single-cells are sorted into plates and lysed. Poly-A mRNA is reverse transcribed with a template switch oligo present. Following exonuclease treatment, splint oligos with 22-RYN UMIs and cellular barcodes are ligated onto the first strand cDNA. Before PCR the libraries are cleaned up. (B) Summary of the datasets generated as part of this study. (C) transcript coverage over reads. Shown are the relative start (grey) and end position (black) for each transcript averaged over the hiPSCs (n=96 cells) (D) Sensitivity to detect RNA molecules in CELLO-seq is shown by summarizing the number of genes detected per error corrected UMI sequence for mouse (n=6; red) and human (n=96; brown). (E) boxplot summarizing the genes per cell (y-axis) with or without exonic heterozygous SNPs probed with CELLO-seq (long) or Smart-seq2 (short) library preparation of hiPSCs (n=96 each), split into genes with at least one heterozygous genic SNP (yellow=allelic) and genes without a heterozygous genic SNP (non-grey=allelic). (F) boxplot of allelic ratio = allele1/(allele1+allele2) (y-axis) depending on exonic heterozygous SNP number (1,2-5, >5) with CELLO-seq (orange) and Smart-seq2 (beige) of hiPSCs. Short read data: 1 SNP (n=6587), 2-5 SNPs (n=10023), > 5 SNPs (n=8088), Long read data: 1 SNP (n=1072), 2-5 SNPS (n=1796), > 5 SNPs (n=1710). (G) boxplot of number of isoforms per cell (y-axis) in mouse 2-cell blastomeres (red) or hiPSCs (brown). Isoforms were grouped in either (all), (known) when they overlapped ENSEMBL transcript ID, (novel), when they did not overlap with an ENSEMBL transcript ID and (TE-derived) when isoforms overlapped repeat and ENSEMBL transcript ID, or when isoforms overlapped with repeats but did not overlap with genic exons. (H) boxplot of mean expression of isoforms (y-axis) by classification of (G) in mouse 2-cell blastomeres (red) or hiPSCs (brown). Mean expression level was calculated from logcounts of all cells. The boxplots shown in E-H show the median, first and third quartiles as a box, and the whiskers indicate the most extreme data point within 1.5 lengths of the box.

## Results

### CELLO-seq produces full-length molecule level cDNAs

CELLO-seq enables single cell analysis by using long read sequencing to profile full-length transcripts (Fig 1A). In brief, we used a 3’-amino blocked template switching oligonucleotide (TSO) to prevent the synthesis of TSO-primed inserts. We also performed the reverse transcription (RT) at high temperature whilst adding RT enhancers to increase cDNA output and the length of transcripts. After RT, we splint-ligated an oligonucleotide containing both a 22-base unique molecular identifier (UMI), and a cellular barcode, to label unique mRNAs. By designing UMIs with a repetitive pattern of RYN (where R and Y represent purine and pyrimidine, respectively), we prevent the presence of homopolymers, which remain inaccurately basecalled in ONT sequencing, although this challenge is receiving continued attention ^22^. Importantly, these long UMIs allow the identification of PCR duplicates, which can be used for deduplication and error correction by producing consensus sequences for each full-length transcript ^23,24^ (Fig 1A). Additionally, and unlike other protocols, we aim for high PCR duplicate numbers by limiting library diversity prior to amplification ^25^ and requiring high cycle numbers to amplify (Fig S1A). These two steps enable us to error-correct transcriptomic libraries sequenced on ONT instruments thus facilitating the generation of high-quality long-reads that can be used for isoform, allelic gene expression analysis and be uniquely mapped back to TEs. However, CELLO-seq was also designed for compatibility with Illumina sequencing, enabling complete transcript coverage.

To assess the performance of CELLO-seq, we generated two independent datasets, the first comprising six manually isolated, 2-cell stage mouse blastomeres which were deeply sequenced, and the second profiling 96 hiPSCs using shallower sequencing (Fig 1B). Confirming the efficacy of our protocol, CELLO-seq produced mostly full-length transcript sequences (Fig 1C): The majority of reads (75%) were flanked with both TSO and oligo-dT-derived sequences, and the mean read length of mapped reads was 2-2.5kb (Fig S1B, D). This contrasts with other recently-published single cell long read protocols, which generated libraries with the 10x protocol before sequencing using ONT or PacBio, where only 40% of reads contained both cellular barcodes and UMIs ^26,27^.

Subsequently, we processed the CELLO-seq libraries using a novel computational pipeline, sarlacc, which demultiplexes cell barcodes, trims adapters, identifies UMIs, and error corrects or deduplicates each cDNA molecule (Fig S1C, https://github.com/MarioniLab/sarlacc). We measured expression for an average of 5,000 genes (200,000 molecules) per cell for the mouse blastomeres and 1,000 genes (10,000 molecules) per cell for the hiPSCs, reflecting the different sequencing depths of these experiments (Fig 1D). Comparing our ONT data with Illumina short read sequencing of the same cDNA, we noted that although the depth of sequencing of the CELLO-seq data is lower when compared to short read sequencing (Fig S1E), ERCC spike-in concentration and read numbers showed good correlation (R=0.84) in both cases, demonstrating that CELLO-seq is quantitative (Fig S1F).

To further demonstrate the utility of CELLO-seq, we examined its ability to quantify allele-specific expression by preparing CELLO-seq and Smart-seq2 libraries ^28^ from cells from two human iPSC lines. Using previously generated phased exome-seq data from the same cell lines ^29^ we were able to measure allelic expression for genes with heterozygous SNPs present in coding regions (Fig 1E, S1G). As expected, the degree of allele-specific expression did not change as a function of the number of exonic SNPs when using the CELLO-seq protocol. By contrast, the Smart-seq2 data showed a clear trend, with more extreme ASE being observed for genes containing only a single exonic SNP (Fig 1F). Consequently, CELLO-seq provides a less biased approach for measuring allele-specific expression than short read strategies.

Next, we probed the ability of CELLO-seq to measure the expression of isoforms in single cells. After processing the error corrected reads with FLAIR ^30^, we were able to measure the expression of ∼10,000 and ∼1,000 distinct isoforms across the mouse blastomeres and human iPSCs respectively, amongst which ∼5,000 and ∼300 were novel (Fig 1G). Amongst these, CELLO-seq enabled detection of 200 and 20 TE-derived isoforms per cell in mouse blastomeres and hiPSCs, respectively (Fig 1G). Interestingly, the expression of TE-derived isoforms in both datasets was higher than all isoforms (Fig 1H), which warrants further investigation. Hence, not only can CELLO-seq detect novel isoforms in single cells but it can also enable study of the role of TE-derived isoforms at single cell resolution.

### CELLO-seq enables analysis of TE-derived isoforms and TEs at unique loci

TEs have long been known to give rise to alternative transcript start and end sites, resulting in the generation of isoforms when the TEs fall near the coding region of existing genes ^31^. Long-read sequencing protocols that enable capture of full-length transcripts, such as CELLO-seq, provide an ideal opportunity to probe their expression. Consistent with previous studies, we found upregulation of TE-derived isoforms, defined as transcripts of non-TE genes whose TSS or TES overlaps a TE, leading to an alternative end or start of the transcript isoform. TE-derived isoforms are clearly visible from ERVL-MaLR elements - ORR1 and MT in mouse blastomeres, and Alu elements in hiPSCs ^4,6,32–34^ (Fig 2, S1H). In the mouse 2-cell blastomeres most TE-derived isoforms stem from SINE B1 and B2 elements (Fig 2A, S1H). DNA transposons also give rise to TE-derived isoforms in both datasets (Fig 2A-D, S1H).

**Figure 2.**
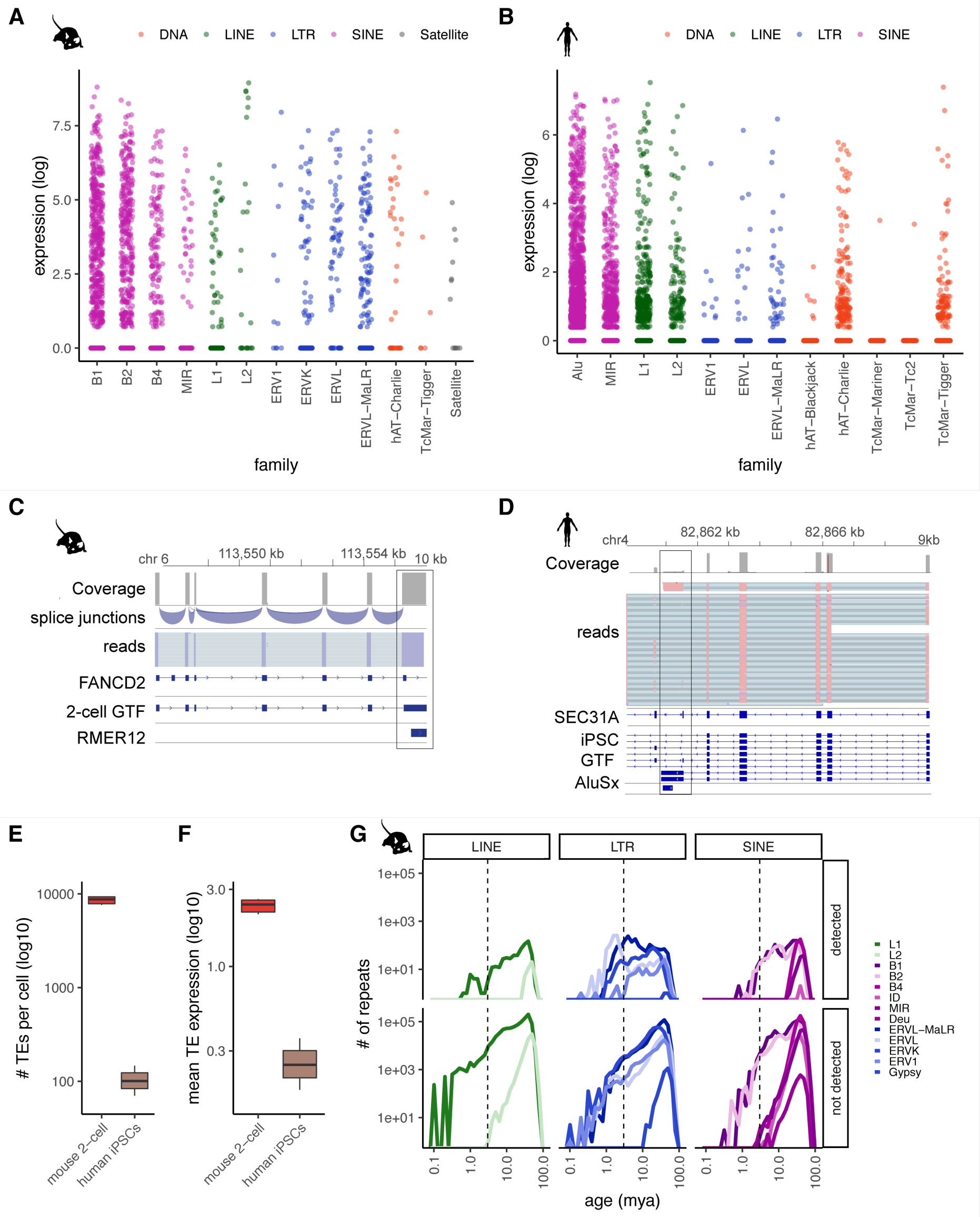
CELLO-seq enables TE-derived isoform and TE expression analysis in single cells at single loci. (A) Jitterplot of expression level (logcounts) of TE-derived isoforms in mouse 2-cell blastomeres stratified by repeat family and coloured by repeat class. B1 (n= 990), B2 (n=630), B4 (n=300), L1 (n=96), L2 (n=24), ERV1 (n=12), ERVK (n=90), ERVL (n=66), ERVL-MaLR (n=174), hAT-Charlie (n=36), TcMar-Tigger (n=6), Satellite (n=12). (B) Jitterplot of expression level (logcounts) of TE-derived isoforms in hiPSCs stratified by repeat family and coloured by repeat class. Alu (n=16643), MIR (n=4628), L1 (n=3649), L2 (n=1602), ERV1 (n=178), ERVL (n=445), ERVL-MaLR (n=534), hAT-Blackjack (n=89), hAT-Charlie (n=2225), TcMar-Mariner (n=89), TcMar-Tc2 (n=89), TcMar-Tigger (n=890). (C) genome browser view of stranded reads overlapping a TE-derived isoform by RMER12 integration in an intron of FANCD2 in mouse 2-cell embryos, exonic reads = blue, introns = turquoise. Shown are all reads from six 2-cell blastomeres. 2-cell GTF has been created by FLAIR using CELLO-seq reads after error correction as input. (D) genome browser view of stranded reads overlapping a TE-derived isoform by AluSx integration into intron of SEC31A in hiPSCs. Shown are all reads from 96 hiPSCs. iPSC GTF has been created by FLAIR using CELLO-seq reads after error correction as input. (E) boxplot of number of autonomous TEs in each cell (y-axis) in mouse 2-cell blastomeres (red) or hiPSCs (brown). (F) boxplot of mean expression of autonomous TEs in each cell (y-axis) in mouse 2-cell blastomeres (red) or hiPSCs (brown). (G) frequency plot of number of repeats (y-axis) by age of TE (mya) (x-axis). Upper panel shows TEs we mapped in our study to mouse 2-cell blastomeres and lower panel shows all repeats from mouse UCSC repeatmasker annotation^35^. TEs are grouped by TE class and coloured by TE family. Repeat class: pink = SINE, green = LINE, blue = LTR, red = DNA, Satellite = grey. The boxplots shown in E and F show the median, first and third quartiles as a box, and the whiskers indicate the most extreme data point within 1.5 lengths of the box.

We then wanted to study the expression of the autonomous TEs themselves. To start, following error correction, we mapped reads to unique TE locations in the genome. In total, we observed expression of an average of 10,000 and 1,000 unique TE loci per cell in mouse blastomeres and hiPSCs, respectively (Fig 2E-G). We observed expression of individual TEs of different ages (see Methods), including those belonging to young (defined as being < 2 mya) subfamilies of TEs. Nevertheless, we noted that very young TEs tended to be less frequently detected in our dataset (Fig 2G, S1I-K). Since such young TE families contain members with highly similar sequences, we hypothesized that this high sequence similarity was impeding our ability to uniquely map reads.

### Simulations show that young human L1 expression can be accurately quantified

To investigate whether this lower detectability was driven by challenges in mapping arising from the high sequence similarity of such elements, we simulated reads in rolling windows from the 3’ end of all young L1PA and L1Md elements for the human and mouse genome, respectively. For each element, we simulated differing read length (1, 2 or 3kb reads - data for 1 and 3kb on github: https://github.com/MarioniLab/long_read_simulations) with either perfect or an ONT read identity error profile (Fig 3F, 1x coverage) ^36^, and considered data with 1x, 5x or 10x coverage of each read (Fig 3A).

**Figure 3.**
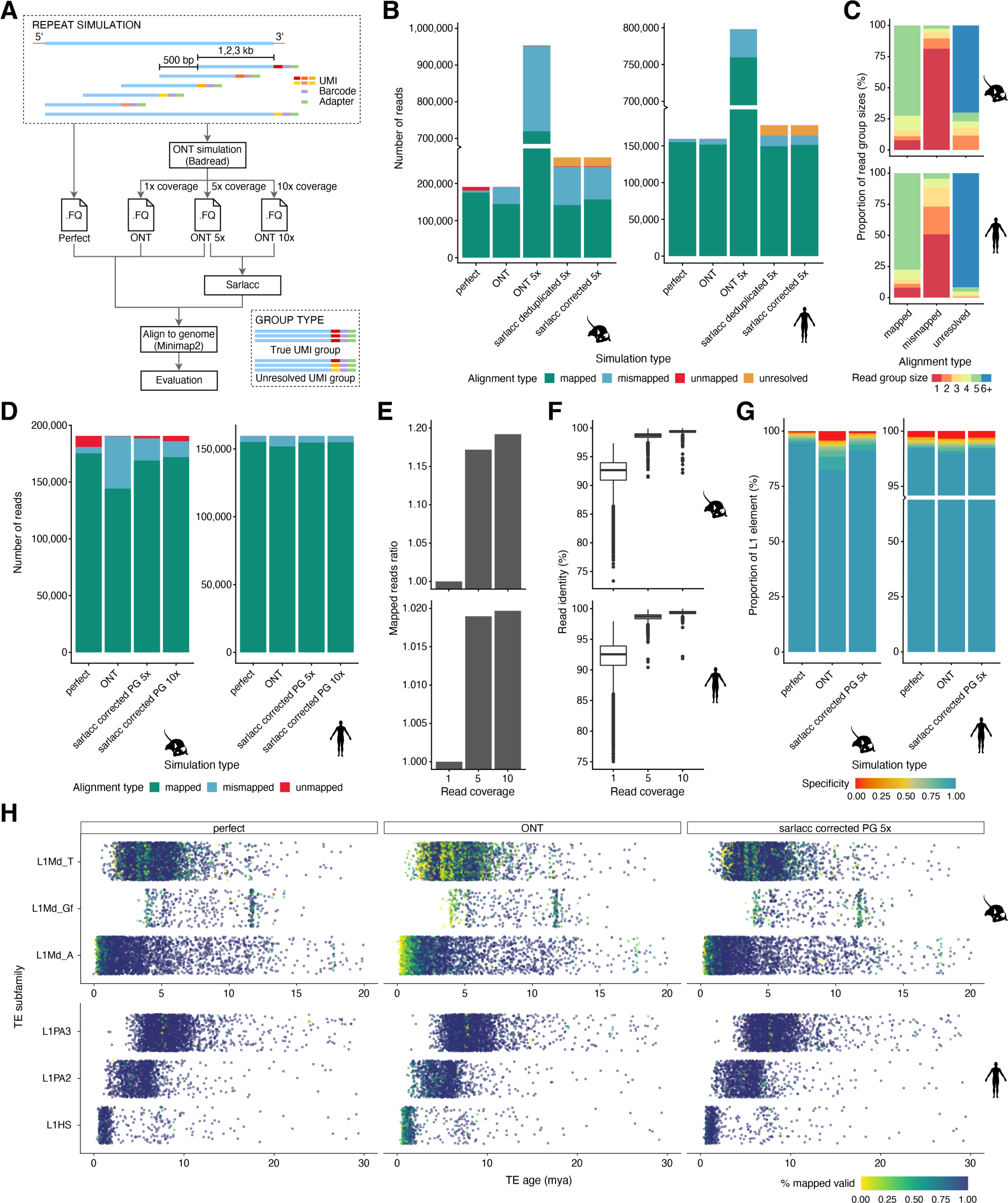
Simulations characterise the mapping of young L1 in mouse and human genome. (A) schematic of strategy of simulation of reads to test mapping of young L1s. Reads were simulated in rolling windows from the 3’ end of all young L1PA and L1Md elements for the human and mouse genome, respectively. For each element, we simulated differing read length (1, 2 or 3kb reads – data for 2kb reads shown) with 500bp gap distance between windows and considered data with 1x, 5x or 10x coverage of each read. Reads were simulated with either perfect sequence identity or with ONT read identity. Reads were mapped either directly to the genome or processed using the sarlacc pipeline to produce both deduplicated and error corrected reads (see methods/Fig S1D). (B) bargraph showing the number of reads (y-axis) by the simulation type (x-axis), colour-coded by alignment type with mapped = turquoise (read at correct location after mapping with minimap2 to the genome), mismapped = light blue (reads mapped at wrong location), unmapped = red (read that could not be mapped with minimap2 to the genome), unresolved = yellow (group with multiple molecules present in one group. Left = mouse L1s, right = human L1s. (C) bargraph of proportion of read group sizes (y-axis) by alignment type (x-axis), colour-coded by group size; Upper panel=mouse data, lower panel = human data. (D) stacked bargraph of number of reads (y-axis) by simulation type, coloured by alignment score as defined for (B). (E) bargraph shows ratio of mapped reads (y-axis) by read coverage (x-axis) for mouse = top panel and human = bottom panel. (F) boxplot of read identity (y-axis) by read coverage (x-axis) for mouse = top panel and human = bottom panel. (G) Stacked bargraph showing proportion of L1 elements (y-axis) by simulation type (x-axis), coloured by specificity score; mouse L1 (left panel) and human L1 (right panel). (H) Jitter plot of TE subfamily (y-axis) by TE age (million years ago) grouped by simulation type and coloured by % mapped reads. Mouse L1 (top panel) and human L1 (bottom panel). Simulation type: perfect = perfect read identity, ONT = ONT read identity, ONT 5x = ONT read identity with 5x coverage of each read, sarlacc corrected 5x = ONT read identity score, 5x coverage run through sarlacc using error correction, sarlacc corrected 10x = ONT read identity score, 10x coverage run through sarlacc using error correction, sarlacc deduplicated 5x = ONT read identity score, 5x coverage run through sarlacc using deduplication by randomly choosing 1 read. PG = perfect grouping. The boxplots shown in F show the median, first and third quartiles as a box, and the whiskers indicate the most extreme data point within 1.5 lengths of the box.

To assess the ability to map reads to the correct locus, we considered three strategies – i) a naïve mapping of reads to the genome; ii) deduplication of reads with identical UMI sequence prior to mapping to the genome; and iii) error correction of reads with identical UMI sequence prior to mapping to the genome (Methods).

As expected, reads with an ONT read identity that were naïvely mapped to the genome were incorrectly mapped more often than perfect reads (Fig 3B; perfect vs ONT). At 5x coverage, a naïve mapping of ONT reads led to a large (∼5 times) increase in the number of mapped reads, artificially inflating the number of correctly and incorrectly mapped molecules. Our bespoke error correction or deduplication strategy substantially removed this artificial inflation of gene counts, but a substantial number of mismapped reads were still observed, especially for the mouse. Additionally, we noted that some grouped sets of reads were unresolved, being comprised of distinct molecules. Finally, we observed that error correction reduced the number of mismapped reads in comparison to deduplication (Fig 3B).

To understand what features might explain the set of reads that could not be properly mapped following error correction, we first compared the distribution of UMI counts in the mapped, mismapped and unresolved groups. For mapped reads, the UMI group sizes were generally equal to the read depth (5x), while mismapped and unresolved reads were grouped into smaller (<5x) and larger (>5x) groups, respectively (Fig 3C). Since this suggested that imperfect grouping of UMI sequences might underpin the erroneously mapped reads, we next asked whether a perfect grouping would improve performance. Consistent with our hypothesis, perfect grouping would allow us to increase the proportion of correctly mapped reads with increased read identity score, especially for the mouse (Fig 3D-F). Finally, we investigated whether we could in principle optimise the grouping by generating reads with longer UMIs (Fig S3).

To this end, we simulated differing UMI lengths (10-50) and differing read depth (5x and 10x) before assessing our ability to correctly group reads based on their UMIs using the Levenshtein Distance (Methods). When simulating data with an ONT error profile, we observed that 50nt UMIs were needed to correctly group the simulated data (Figure S3B-C). This contrasts with the relatively short (∼10nt) UMI used for NGS and the 22nt UMI used in CELLO-Seq.

In sum, for the large majority of L1s in both human and mouse we determined that CELLO-seq reads could be correctly mapped, with the exception being very young mouse L1s due to difficulties in grouping reads (Fig 3H). To avoid these loci confounding downstream analyses of our CELLO-seq data, for each L1 element we calculated a specificity score based on the number of mapped reads from the selected L1 element and the number of mismapped reads from other L1 element that aligned within the location of the selected L1 element (Table on GitHub: https://github.com/MarioniLab/long_read_simulations): 98% of the human L1s have a specificity score > 98% (with an increase to 98.5% for perfect sequence complementarity), while for mouse L1s, 80% of L1s have a specificity score > 80% with our current method (Fig 3G). Henceforth, we consider filtered datasets, only including L1s with a specificity score of 80% (Table S1).

### Studying the locus-specific expression of TEs reveals that subfamilies react differently to global epigenetic reprogramming

During preimplantation development the whole genome is epigenetically reprogrammed, which in turn leads to substantial changes in the transcriptome. Consistent with previous studies, we observe a genome-wide increase of TE expression in the sequenced mouse blastomeres at 2-cell stage embryos (Fig 4A, S4)^4,32^. Interestingly, in contrast to SINE and MERVL elements, young full-length LINEs were consistently expressed from only a few loci in all of the 2-cell blastomeres (Fig 4A-C, S4A, B).

**Figure 4.**
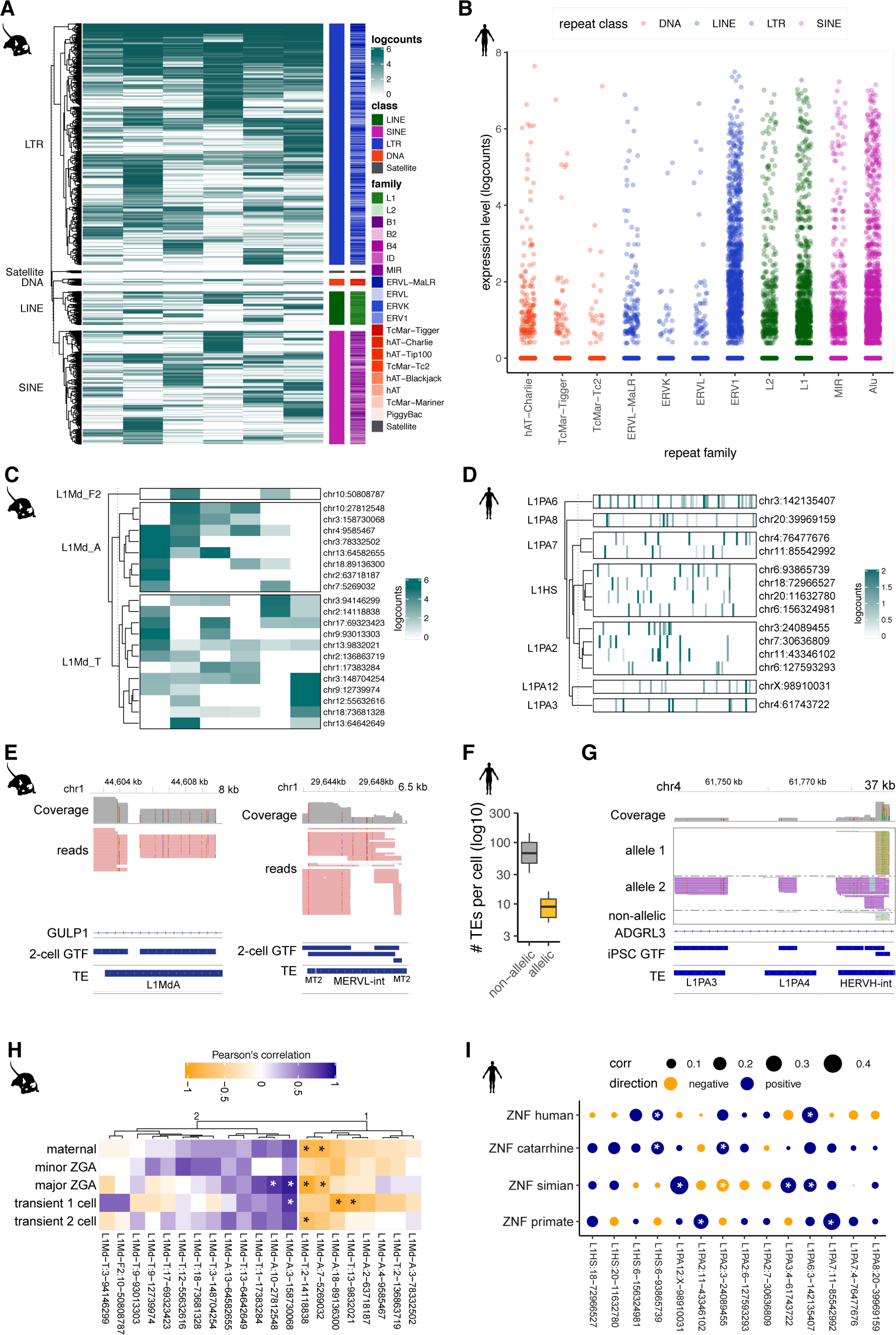
CELLO-seq enables study of young TEs at unique loci. (A) Expression of all highly expressed TEs (n=3618) (mean expression across all cells > 1 logcounts) in 2-cell mouse embryos. Clustering is performed by TE class. B1 (n=450), B2 (n=380), B4 (n=167), ERV1 (n=65), ERVK (n=282), ERVL (n=694), ERVL-MaLR (n=1159), hAT (n=1), hAT-Blackjack (n =2), hAT-Charlie (n=38), hAT-Tip100 (n=1), ID (n=2), L1 (n=285), L2 (n=19), MIR (n=35), PiggyBac (n=1), Satellite (n=20), TcMar-Mariner (n=1), TcMar-Tc2 (n=1), TcMar-Tigger (n=15). (B) Expression of all long (> 5 Kb) L1 elements that are highly expressed (mean >1 logcounts) in mouse 2-cell blastomeres clustered by their subfamilies. (C) genome browser view of stranded reads overlapping L1MdA (left) and MERVL-int (right) in mouse 2-cell embryos, sense reads = red. Shown are all reads from 6 2-cell blastomeres. 2-cell GTF has been created by FLAIR using CELLO-seq reads after error correction as input. (D) Pearson correlation of preimplantation networks ^38^ with unique loci of young, long (>5.8 kb) L1 elements in mouse 2-cell blastomeres as a heatmap. (E) Expression of all TEs (n=414) in hiPSCs shown by family and coloured by TE class (sum of expression across all cells > 10 logcounts). hAT-Charlie (n=1335), TcMar-Tigger (n=534), TcMar-Tc2 (n=178), ERVL-MaLR (n=1157), ERVK (n=178), ERVL (n=534), ERV1 (n=3738), L2 (n=2937), L1 (n=9078), MIR (n=3827), Alu (n=12905). (F) Expression of all long (> 5 Kb) TE elements that are highly expressed (mean >1 logcounts) in hiPSCs clustered by their subfamilies. (G) boxplot of mean number of TEs per cell (y-axis) which we can call allelically (yellow) or non-allelically (grey). The boxplots show the median, first and third quartiles as a box, and the whiskers indicate the most extreme data point within 1.5 lengths of the box. (H) genome browser view of stranded reads overlapping L1PA3, L1PA6 and HERVN-int in hiPSCs. Shown are all reads from 96 hiPSCs. iPSC GTF has been created by FLAIR using CELLO-seq reads after error correction as input. yellow = allele1 reads, purple = allele2 reads, grey = non-allelic reads. (I) Pearson correlation of ZNFs ^39,40^ with unique human full-length TEs (>5.8kb) with negative correlation shown in blue and positive correlation shown in yellow and the size of the dot showing the correlation coefficient as a bubble graph. Significant associations (FDR < 0.1 = *).

Given the ability of CELLO-seq to detect locus specific TE expression, we had the opportunity, for the first time, to explore whether the expression of specific TEs is associated with the expression of different classes of protein coding genes and to infer putative regulatory mechanisms. Prolonged L1 expression is associated with the arrest of mouse preimplantation development ^20,21^. CELLO-seq allowed us to test the correlation between the expression of genes specifically associated with the maternal, zygotic, 1 and 2-cell stages of development and the expression of young L1s in mouse 2-cell blastomeres. Consistent with previous work, the majority of young L1 loci were positively correlated with gene networks associated with different stages of preimplantation development. Interestingly, we also observed a subset of L1 loci with a statistically significant negative correlation with early development expression profiles, suggesting that these TEs might be important for the transition between 2-cell and gastrulation stages (Fig 4D). To test the biological relevance of this finding, a larger sample size and follow up experiments are required in order to investigate whether knockdown of specific L1s can lead to disruption of preimplantation development ^20,21^.

CELLO-seq also enabled us to measure expression at unique young L1, Alu, and HERV-int elements in hiPSCs (Fig 4E, S4C). Coherent with previous findings in bulk and single cell NGS datasets ^6,33,34^ Alu elements contributed the highest number of expressed loci, while the most highly expressed TEs were HERV-int elements and their LTRs – LTR7 (Fig 4E, S4C). CELLO-seq permitted us to detect locus-specific expression of four L1Hs and four L1PA2s (Fig 4F,G). Additionally, we were able to study allelic TE expression at single cell resolution, with a mean number of ten TEs showing allele-specific expression in each hiPSC (Fig 4H). This is an important property of CELLO-seq as polymorphic, heterozygous integrations of TEs have been widely observed in humans ^37^.

Next, we focused on the correlation of transcriptional repression of TEs through known epigenetic regulation of KRAB-ZFPs ^39,40^. KRAB-ZFPs are characterised by a tandem array of zinc finger nucleases (ZNFs) and a KRAB domain that recruits KAP1, which functions as a scaffold for other repressive histone-modifiers and binding factors. The expression of TEs is in an evolutionary arms race with the transcriptional activity of their sequence specific ZNFs ^39,40^. Our locus specific hiPSC data allowed us to examine this relationship in more detail. Overlaying the expression of classes of ZNFs that have evolved at different time points during primate evolution with full-length TEs expressed in hiPSCs, we generally detected a (significant) positive correlation (Fig 4I). Furthermore, two L1 loci (L1PA6:3-142135407 and L1HS:6-938665739) show statistically significant positive correlation with young human ZNFs, suggesting that these L1s have potentially escaped ZNF transcriptional control via specific mutations in their ZNF binding sites. This shows that locus specific correlation of expressed TEs and their transcriptional regulators enables discovery of putative regulatory mechanisms and that previous studies, which aggregated data across all L1Md subfamilies, do not have sufficient granular resolution to interrogate how the expression of individual TEs might be regulated.

## Discussion

CELLO-seq represents a powerful new tool for profiling the expression of full-length molecules, including transposable elements, in single cells. Unlike other approaches that have generated long transcript reads from single-cells, CELLO-Seq does not rely on library generation using the 10X Chromium, or other similar platforms, but rather generates libraries that can be immediately sequenced using ONT. As a result, in the development of CELLO-seq we have been able to optimise conditions for RT and PCR, and to link our oligonucleotide design to our bespoke analysis pipeline, all of which enable full-length molecules to be captured and quantified more reliably. A crucial feature of CELLO-seq is the ability to take advantage of PCR duplicates to error correct reads prior to mapping. Moving forward, we envisage that the efficiency of this error correction step could be improved by increasing the number of PCR duplicates for each read via target enrichment, or by performing emulsion PCR to inhibit the PCR jackpot effect ^41^, thereby flattening the distribution of PCR duplicates among UMIs (Figure S1A).

Even with the innovations of CELLO-seq, challenges remain. For example, measuring expression at very young TEs is challenging due to their very high sequence similarity. Indeed, using simulations, we demonstrated that while we could map L1s across the human genome with high specificity, mapping young mouse L1s to unique loci with high specificity is currently only possible for ∼85% of loci. To study TEs with very high sequence identity, our simulations suggest that we will need to increase the UMI length in CELLO-seq to 50nt in order to enable accurate grouping of reads with ONT read identity error profile. We note that while a 50nt UMI is much longer than any used by existing single-cell technology, it would be feasible in CELLO-seq since we use a splint-ligation to add the UMI sequence. ONT sequencing of rolling circle amplification products has also been demonstrated as a means of delivering error-corrected transcript reads ^42,43^. However, the raw read throughput of these protocols is low compared to CELLO-seq, and the number of independent observations of each transcript remains highly left-skewed, with ∼99% of reads containing fewer than 4 concatemers, constituting an inherent ceiling to error-correction fidelity that cannot be improved by increasing sequencing depth. In contrast, PacBio sequencing, which also exploits rolling-circle replication, can now deliver very high fidelity reads (∼Q30) due to the many polymerase passes of each insert seen in recent chemistry upgrades. However, the unique read throughput remains constrained by the fixed number of zero mode waveguides manufactured on each flow cell, rendering this platform relatively expensive.

From a biological perspective, our data already demonstrate how CELLO-seq can be used to provide insight into the transcription and regulation of TEs. We have shown that expression of TEs is heterogeneous across members of a TE subfamily and, moreover, that members of the same subfamily are differentially correlated with the expression of putative regulatory elements. This presents the intriguing possibility that different classes of epigenetic regulators can impact the expression of unique TEs from the same family, potentially leading to stage-specific expression during development.

In conclusion, CELLO-seq enables isoform, allelic and TE expression at unique loci at single cell resolution. Our simulations show that we are able to map autonomous TEs with high specificity at most loci. The possibilities to study the role of transposable elements at unique loci – the same way we study protein coding genes - will provide novel insight into TE biology and the role of TEs in gene expression.

## Material and Methods

### Cell culture of human-induced pluripotent stem cells

Human-induced pluripotent stem cells (hiPSCs) were thawed and cultured under feeder-free conditions on Vitronectin XF™-coated (Stem Cell, #07180) 10 cm tissue culture treated plates (Corning) in complete Essential 8 medium (Life Technologies #A1517001) supplemented with 1% Penicillin/Streptomycin (Invitrogen, #15140122). Cells were incubated at 37°C, 5% CO2 and media changed every day except for the day of passaging. Cells were routinely passaged using 0.5 mM EDTA (Life Technologies #AM9260G) at least 3 times post-thawing before collection. HiPSCs were harvested using Accutase (Millipore, #SCR005) to generate a single cell suspension. Single cells were resuspended in 1X PBS (Thermo Fisher Scientific, #10010023) and passed through a 40uM filter (Corning #CLS431750-50EA) before FAC sorting (BD Influx™ Cell Sorter) into 96-well plates (Framestar, #4TI-0960) containing 2µl lysis buffer with 1U/µl Rnase Inhibitor (RRI) and ERCC spike in mix at a 1:20 dilution (Ambion Cat #4456740). Plates were promptly sealed (Thermo Fisher Scientific #AB0626), spun down and frozen at −80 °C until further processing.

### Collection of mouse preimplantation embryos

#### Ethical considerations

All experimental procedures were in accordance with UK Home Office regulations and the Animals (Scientific Procedures) Act 1986 (PPL No: P418B15F6). All experimental protocols were approved by the Animal Welfare and Ethical Review Body (AWERB) of the University of Cambridge CRUK Cambridge Institute. At the end of the study, mice were euthanized by cervical dislocation, in accordance with the above stated UK Home Office regulations.

#### Reagents

Human chorionic gonadotropin (Chorulon; National Veterinary Services, #804745), Pregnant mare serum gonadotropin (PMSG; National Veterinary Services, #859448), EmbryoMax KSOM medium with 1/2 amino acids without BSA (Reagent setup; Merck Millipore, cat. no. MR-107-D), BSA powder (Sigma-Aldrich, cat. no. A3311), M2 medium with HEPES (Sigma-Aldrich, cat. no. M7167), Cytochalasin B (Reagent setup; Sigma-Aldrich, #C6762), Tyrode’s solution, acidic (Sigma-Aldrich, #T1788), Dimethyl sulfoxide (DMSO; Sigma-Aldrich, #D2438). Supplemented EmbryoMax KSOM medium: Before separation of blastomeres from embryos, further supplement 50.0 mL of EmbryoMax KSOM medium with 20.0 μL of phenol red solution (0.5% (wt/vol) in H2O) and 3.0 mg/mL BSA. The medium should be prepared in a sterile tissue culture hood and can be stored at 4 °C for 2 weeks. Warm the medium in a 37 °C, 5% CO2 incubator for at least 30 min before use.

Cytochalasin B stock solution: Prepare a stock solution of 1.0 mg/mL cytochalasin B by resuspending 1.0 mg of cytochalasin B in 1.0 mL of DMSO. Divide the solution into single-use aliquots, 7.5 µL each, and store at −20 °C for up to 6 months.

#### Collection of blastomeres from embryos

To collect blastomeres from two-cell stage mouse embryos, female C57BL/6J mice older than 8 weeks of age were super-ovulated by intraperitoneal injection of pregnant mare serum gonadotropin (7.5 IU per mouse) at 4PM. 46–48 h later (i.e., 2–4 pm), each mouse was injected with 7.5 IU of human chorionic gonadotropin and left to mate with proven studs (C57BL/6J). Vaginal plugs were checked the following morning (0.5 d.p.c.) and mice that have successfully mated were used for dissection.

For late 2-cell stage embryos collection, the plugged mouse was culled at 4:30 pm, and the uterine horns and oviducts of the donor mouse were dissected under a dissection microscope and placed on the lid of a 100-mm tissue culture dish in M2 medium. The oviduct with part of the uterine horn was cut out of the uterus and place it into M2 medium. Under a dissection microscope, a needle was inserted infundibulum and the oviduct was flushed with M2 medium, with late 2-cell stage embryos being flushed out into a dish. The embryos were cultured in 1.0 mL of supplemented EmbryoMax KSOM medium with phenol red (containing 3.0 mg/mL BSA) in a 5% CO2 humidified incubator at 37 °C. Just prior to separating the blastomeres, embryos were cultured in 1.0 ml of supplemented EmbryoMax KSOM medium containing 3.0 mg/mL BSA and 7.5 μg/mL cytochalasin B for at least 30 min. A dish with four drops of M2 medium with 7.5 μg/mL cytochalasin B and two drops of Tyrode’s solution was prepared. Under a dissection microscope, the embryos were transferred to M2 medium with 7.5μg/mL cytochalasin B. All embryos were transferred into the first drop of Tyrode’s solution (washing step) and then into the second drop of Tyrode’s solution. We waited until the zona pellucida of the embryos had dissolved, and quickly transferred the embryos into the first M2 drop, and washed the embryos through the second and third M2 drops. The blastomeres were transferred to an M2 drop immediately to minimize the damage of the acidic Tyrode’s solution to blastomeres. A thin capillary pipette with a blunt end was used to aspirate and blow out the blastomeres a few times to dissociate them. The individual blastomeres were then transferred into the fourth M2 drop, where they remained (for up to 30 min) prior until further processing.

Single cells were subsequently mouth pipetted into 96-well plates (Framestars #4TI-0960) containing 2 µL lysis buffer with 1 U/µL Rnase Inhibitor (RRI), ERCC spike in mix at 1 in 20Mio dilution (Ambion #4456740). Plates were sealed (Thermo Fisher Scientific #AB0626) immediately spun down and frozen at −80 °C until further processing.

### CELLO-seq

Before starting with reverse transcription, we first eliminated any secondary structure in the RNA. We added a 2.2 µl mastermix of 1 µl (1mM) dT-oligo (IDT), 1 µl (1µM) dNTPs (25 mM each, Thermo Fisher Scientific #R0182) as well as 0.2 µl (20U/µL) of RNAse inhibitor (Superase In, Thermo Fisher Scientific, #AM2694) and incubated the mix at 72°C for 3min before returning to ice. The reverse transcription was performed in 10 µl preparing a 5.8 µl mastermix with 1 µl of 200U/µl Superscript IV Reverse Transcriptase (Invitrogen #18090050), 2 µl of 5x Superscript IV RT buffer, 1 µl of 5M betaine, 0.1 µl of (100µM) template switch oligo (IDT), 0.5 µl of 100mM DTT, 0.1 µl ET SSB (NEB #M2401S) and 1.6 µl nuclease free H2O (DNase/RNase-Free Distilled H2O, Thermo Fisher Scientific #10977-049). cDNA synthesis and template switching were performed for 10 min at 57 °C and 120min at 42 °C. After RT we digested all residual oligos using adding 5 µl mastermix of 4 µl nuclease free H2O (DNase/RNase-Free Distilled H2O, Thermo Fisher Scientific #10977-049), 0.5 µl of Exonuclease I (NEB #M0568) in 0.5 µl of 10x Cutsmart buffer (NEB # B7204S). We incubated the reaction at 37 °C for 1h, followed by 95 °C for 3 min to fully denature both the exonuclease and cDNAs. Before performing the splint ligation, we annealed the splint oligos in 1 µl of 10x Oligo hybridisation buffer (500 mM NaCl, 10 mM Tris-Cl, pH 8.0, 1 mM EDTA, pH 8.0), 1 µl of 100 µM top splint (IDT) and 1 µl of 100 µM bottom splint (IDT) diluted in 7 µl nuclease free H2O (DNase/RNase-Free Distilled H2O, Thermo Fisher Scientific #10977-049). The reaction was annealed at 95 °C for 1m following 20 s at 95 °C and decreasing temperature by 1 °C every cycle, this step was repeated 80 times and subsequently kept at 4 °C until further processing.

The splint ligation reaction was performed by preparing a 5 µl mastermix that consisted of 0.5 µL Hifi Taq Ligase (NEB #M0647S), 0.5 µl 10x Taq ligase buffer, 1µl of 10 µM annealed splint oligo, 0.25 µL NAD+ (NEB #B9007S) and 2.75 µl nuclease free H2O (DNase/RNase-Free Distilled H2O, Thermo Fisher Scientific #10977-049). The ligation was performed at 55 °C for 1h. Before clean-up we performed a Proteinase K digestion, as the beads were sticky with leftover protein. We digested the proteins with Proteinase K (NEB #P8107S) for 5 min at 50 °C. Barcoded cDNA was then pooled in 2 mL DNA LoBind tubes (Eppendorf #0030108051) and cleaned up using AMPure XP beads at a ratio of 0.5X volume. Purified cDNA was eluted in 12.75 µl H2O. 0.75 µl of the purified DNA was used to perform a proxy qPCR to avoid over-amplification. For the qPCR we used a 25 µL reaction with 12.5µl of 2x KAPA HiFi Uracil+ hot start polymerase mix (Roche # KK2800), 1.25µl of 10 µM forward and reverse PCR primers (IDT), 5µl of 5M betaine, 0.5 µL ET SSB, and 1.25µl of 2x EvaGreen (Biotium #31000-T). The qPCR was performed for 3 min at 90 °C for initial denaturation followed by 29 cycles of 20 s at 90 °C, 30 s at 65 °C, 10 min at 72 °C for 40 cycles. We then used four times fewer cycles than indicated by the qPCR to lead to logarithmic phase of amplification and prepared a PCR reaction of 50 µl PCR master mix consisting of 25µl of 2x KAPA HiFi Uracil+ hot start polymerase (Roche # KK2800) mix, 2.5µl of 10 µM forward and reverse PCR primer, 10µl of 5M betaine, 0.5 µL ET SSB. The PCR was cycled as given: 3 min at 90 °C for initial denaturation followed by 29 cycles of 30 s at 90 °C, 15 s at 65 °C, 10 min at 72 °C, for 25-29 cycles, depending on the qPCR results. These high cycle counts are required to yield the high cDNA inputs (>500 ng) required for optimal ONT sequencing. Final elongation was performed for 10 min at 72 °C. Following amplification, all samples were purified using AmpureXP beads at a volumetric ratio of 0.5X with a final elution in 13 µl of H2O (DNase/RNase-Free Distilled H2O, Thermo Fisher Scientific #10977-049). The cDNA was then quantified using the Qubit High Sensitivity dsDNA Assay Kit (Thermo Fisher Scientific, #Q32851). Size distributions were checked on High-Sensitivity DNA chips (Agilent 2100 Bioanalyzer, #5067-4626). All oligo sequences can be found in Table S2.

### Smart-seq2

We prepared cDNA of human iPSCs following the Smart-seq2 protocol ^44^.

### Library preparation and sequencing

#### Short read library prep from CELLO-seq derived cDNAs

For tagmentation we used custom made Tn5 provided by the Protein Expression and Purification Core Facility of the European Molecular Biology Laboratory (EMBL). We subsequently largely followed the previously published protocol with small adjustments^45^. We first annealed the oligos in Oligo hybridisation buffer (500 mM NaCl, 10 mM Tris-Cl, pH 8.0, 1 mM EDTA, pH 8.0), and we used Tn5MErev ^46^ and Nextera R1 oligo (IDT, Table S2) at a final concentration of 350 µM in total 5.15µl H2O. The reaction was annealed at 95 °C for 1 min following 20 s at 95 °C and decreasing temperature by 1 °C every cycle; this step was repeated 80 times and the resulting material kept at 4 °C until further processing. We then loaded the annealed oligo onto the Tn5 using 35 µM annealed oligo in 10µl Tn5 with 0.5µl Oligo Hybridisation buffer (500 mM NaCl, 10 mM Tris-Cl, pH 8.0, 1 mM EDTA, pH 8.0). We incubated for 30min at 23 °C. The tagmentation was performed in 10 mM Tris-HCl pH 7.5 (Merck, #93363), 10 mM MgCl2 (Merck, #M1028), and 25% dimethylformamide (DMF) (Merck, # 227056) using 200 pg target DNA and incubated at 55 °C for 3 min in a preheated thermocycler. To unbind the Tn5 we performed a proteinase K digestion using 0.5 µl proteinase K in 1µl yeast tRNA in 10 mM Tris-HCl pH 7.5 and incubated for 5min at 55 °C. The product was purified using SPRI beads at a ratio of 1:1. We eluted in 12.75µl H2O and used 0.75µl to perform a proxy qPCR in order to not overcycle. The qPCR was performed as above using KAPA HiFi hot start polymerase and Illumina i5 and i7 primers (Table S2). We included a 3 min 72 °C step before starting the initial denaturation, annealed at 67 °C and performed the elongation for 30 s instead of 10 min. We then used 6 times less cycles than the qPCR indicated to perform the PCR as stated above using KAPA HiFi hot start polymerase and Illumina i5 and i7 primers. We included a 3 min 72 °C step before starting the initial denaturation, annealed at 67 °C, performed the elongation for 30 s instead of 10 min and performed 11 PCR cycles. The product was bead purified at 0.6X. The libraries were checked on an Agilent 2100 Bioanalyzer and quantified using Qubit as well as a KAPA library quantification kit, as per manufacturer’s recommendations. Libraries were paired-end sequenced on an Illumina HiSeq 2500 instrument.

#### Short reads for Smart-seq2

Amplified cDNA produced by Smart-seq2 was, in lieu of the Nextera XT tagmentation recommended in the protocol as written, fragmented and processed into an Illumina sequencing library using the NEB Ultra II FS kit (NEB cat. no. E7805S). Libraries were paired-end sequenced on an Illumina HiSeq 4000 instrument.

#### Long reads

200 fmol of CELLO-seq libraries were prepared for ONT sequencing using the Ligation Sequencing Kit (ONT SQK-LSK109) and sequenced using one MinION R9.4.1 flow cell.

### Primary data processing

All commands and parameters for running the software are defined in the associated GitHub repository (https://github.com/MarioniLab/CELLOseq).

#### Short reads

The human iPSCs Smart-seq2 fastq data were processed with FilterByTile from the BBMap (38.76) ^47^ package, Trim Galore (0.6.5) ^48^ and GATK (4.1.4.0) ^49^ for filtering of low quality reads, adapter removal and deduplication, while the mouse 2-cell blastomeres fastq data were processed using UMItools (1.0.0) for demultiplexing and deduplication. STAR (2.7.3a) was then used for alignment to efficiently generate expression profiles for barcoded UMI data ^50,51^. For human iPSCs, we mapped to the human reference genome (gencode, GRCh38_p13) while mouse cells were mapped to the mouse genome (gencode, GRCm38_p6) concatenated with the ERCC reference.

We used featureCounts (2.0.0) for genic count data from the reads, with gencode GRCh38_p13 for human and GRCm38_p6 for mouse ^52^01/10/2020 11:01:00. After initial data processing, we filtered cells by spike-in normalisation, total library counts and mitochondrial RNA contamination following the Orchestrating Single-Cell Analysis with Bioconductor pipeline ^53^.

#### Long reads

After MinION sequencing, we basecalled the raw ONT reads using the high-accuracy model implemented in Guppy (3.5.2) and then preprocessed the reads using the sarlacc pipeline (Fig S1D). We first checked for internal adapters, which could arise from blunt ligation during ONT library preparation, and used Porechop (0.2.4) ^54^ to split the reads if more than 20% of reads had internal adapters. Otherwise we filtered out reads with internal adapters present. We then performed adapter quality control, where we aligned the 250 bp sequence on either end of the read against dT-oligo and TSO sequences to identify the adapters at either end of the full-length cDNA, followed by filtering for reads with both adapters present^55^. We demultiplexed the samples based on the sample barcodes by grouping barcodes that have a Levenshtein distance below the grouping threshold. To reduce the search space for the UMI grouping, we performed pre-grouping by mapping the reads to the relevant transcriptome (defined above), ERCC spike-ins as well as unique genome repeat locations of mouse and human UCSC RepeatMasker track with minimap2 (2.17-r954-dirty) ^55^. We grouped the reads by their UMI sequence within each pregroup and either error corrected the reads by taking the consensus sequence from multiple sequence alignment of up to 50 reads in the UMI group, or by picking a random read from the UMI group in deduplication mode. For this study we used error corrected reads and aligned them back to the genome with minimap2.

#### Allelic gene expression analysis

We lifted over the phased vcf files from the Hipsci consortium using liftOver from UCSC. We then aligned the corrected reads against the genome using minimap2 and ran whatshap (0.18) to phase the reads^56^. The phased bam files were used in the isoform analysis.

#### Isoform expression analysis

To acquire splice-junction corrected isoform data, we corrected the reads using FLAIR (Full-Length Alternative Isoform analysis of RNA, 1.4.0) ^30^. The reads were first mapped to the transcriptome with repeat sequences and ERCCs added using flair-align. We then run flair-correct to perform splice-junction correction by using short read data for the human and mouse genome. During flair-collapse, the reads are grouped by their isoform and are corrected within each isoform separately. Afterwards, we used flair-quantify with salmon to generate isoform count data for each cell^57^. We defined known isoforms as those with known ensembl transcript ID. Any isoforms lacking a known transcript ID were further subdivided into those overlapping repeats, termed TE-derived isoforms, or those that did not overlap a RepeatMasker entry, termed novel isoforms. To define TE-derived isoforms, we filtered the flair derived isoform GTF file using bedtools (2.29.2) by considering only those isoforms whose transcription start site or transcription end site fell within a repetitive element, defined using the RepeatMasker annotation and the nested repeat file downloaded from UCSC.

#### Transposable element expression analysis

For autonomous transposable element analysis, we downloaded RepeatMasker ^35^ and nested repeat annotation from UCSC in GTF format. For the final flair alignment, we used the transcriptome and repeat GTF to quantify TE-derived isoforms as well autonomous TEs.

#### Read normalisation

We used the isoform count data from flair-quantify and then filtered cells by spike-in normalisation, total library counts and mitochondrial RNA contamination following the Orchestrating Single-Cell Analysis with bioconductor (3.10) pipeline in R (3.6.3).

#### Transposable element age analysis

The age of transposable elements was extracted by converting the milliDivergence of each transposable element from the RepeatMasker annotation ^35^ into million years of age using the Jukes-Cantor model.

### Repeat simulation

For repeat simulation, we obtained repeat sequences of all young L1PA and L1Md elements with length >= 2kb from the UCSC repeatmasker track. We then generated simulated reads using a rolling window of size 1, 2 and 3 kb, and gap size of 500 bp starting from the 3’ end of the L1 elements to better simulate the type of reads captured by CELLO-seq. For perfect simulation, we considered Illumina reads as our perfect baseline and therefore assigned Illumina high quality score (Q40) for each of the simulated reads, while for ONT simulation, we utilised a modified version of Badread to simulate ONT read identity and read coverage of 1x, 5x or 10x for each of the simulated reads. The identity parameter used for Badread simulation is 92,96,2.5 – corresponding to a mean read identity of 92%, maximum read identity of 96% and standard deviation of 2.5% – based on the read identity distribution of our ONT reads. We also used a custom error model and qscore model for Badread simulation, which are both trained from our ONT data. To allow for deduplication and error correction of the ONT simulated reads, we included the 3’ adapter utilised by CELLO-seq containing a shared cell barcode and unique simulated 22 bp RYN UMI for each of the simulated reads.

To assess the ability to map the simulated reads to their valid loci, we first processed the ONT simulated reads with 5x and 10x coverage using the sarlacc pipeline in order to produce both deduplicated and error corrected reads. We modified the sarlacc pipeline for the repeat simulation by aligning the simulated reads to the repeat sequences of the young L1PA and L1Md elements in the alignment stage. We then aligned the perfect simulated, ONT simulated and sarlacc processed reads against the reference genome using minimap2 and evaluated the location of the aligned reads against the true location of the simulated reads, which are stored in the simulated reads’ name.

We classified the reads into three categories based on the alignment type - mapped, for reads whose alignment location overlaps the true location by at least 1 bp; mismapped, for reads whose alignment location does not overlap the true location; and unmapped, for reads which did not get aligned. We also added an additional category for sarlacc processed reads - unresolved - for read groups formed by the sarlacc pipeline that are composed of multiple UMIs. We then summarise the information from each read at the L1 element level to calculate the fraction of reads that are mapped, mismapped and unmapped. We also calculated the specificity score for each L1 element based on the ratio of mapped reads from the selected L1 element and the number of mismapped reads from other L1 element that aligned wrongly within the location of the selected L1 element.

For read identity calculation, we ran minimap2 with -c flag in order to generate a CIGAR string for each alignment. The read identity for each read is then calculated based on the number of bases from the read sequence that matches the reference sequence and the number of bases from the read sequence that either does not match (mismatch) or is missing from the reference sequence (insertion/deletion).

We have published the pipeline used for repeat simulation, alongside the result of the young L1PA and L1Md element simulation for 1 and 3 kb reads, in GitHub (https://github.com/MarioniLab/long_read_simulations) to allow for simulation evaluation for other repeat types.

### UMI simulation

For UMI simulation, we generated 10,000 UMI-only reads of length 10, 20, 30, 40 and 50 bp with RYN or NNN pattern. We then performed perfect simulation of the UMI reads with perfect read identity score at a read coverage of 1x, 5x and 10x, as well as ONT simulation using Badread to generate UMI reads with ONT read identity and read coverage of 1x, 5x and 10x, as described previously (data for 1x and 10x read coverage on Github, https://github.com/MarioniLab/long_read_simulations). Levenshtein distances were calculated across all true UMI pairs of perfect simulated UMI with 1x read coverage using the expectedDist function of the sarlacc R package. Group size and group purity evaluation were performed on the UMI groups produced by the umiGroups function of the sarlacc R package with the Levenshtein distance threshold for grouping set to 2, 6, 10 and 14. For no pregrouping evaluation, no pre-grouping of UMI were provided to the umiGroups function, while in pregrouping evaluation, the UMIs were first pre-assigned into groups of 100 unique UMI based on the true UMI sequences and this pre-grouping information was provided to the umiGroups function.

## Abbreviations

(CELLO-seq): CELl LOng read RNA sequencing
(PacBio): Pacific Biosciences
(MaLRs): Mammalian apparent LTR-retrotransposons
(SNP): single nucleotide polymorphism
(sc): single cell
(TE): Transposable element
(ERV): endogenous retrovirus
(LINE): Long interspersed element
(SINE): Short interspersed element
(RNAseq): RNA sequencing
(hiPSCs): Human induced pluripotent stem cells
(RT): Reverse Transcription
(LTRs): long terminal repeat elements
(NGS): next generation sequencing
(ONT): Oxford Nanopore technologies
(UMIs): unique molecular identifiers
(TSO): template switch oligo
(ZNFs): Zinc finger nucleases
(nt): Nucleotides
(bp): Base pairs
(TSS): Transcription start site
(TES): Transcription end site
(ERCC): External RNA Controls Consortium

## Declarations

### Ethics approval and consent to participate

All experimental procedures were in accordance with UK Home Office regulations and the Animals (Scientific Procedures) Act 1986 (PPL No: P418B15F6). All experimental protocols were approved by the Animal Welfare and Ethical Review Body (AWERB) of the University of Cambridge CRUK Cambridge Institute.

### Availability of data and materials

The datasets generated and/or analysed during the current study are available under ArrayExpress accession E-MTAB-9577.

For data analysis the code is available in the following GitHub repositories: https://github.com/MarioniLab/CELLOseq, https://github.com/MarioniLab/sarlacc, https://github.com/MarioniLab/long_read_simulations.

### Authors’ contributions

RVB conceived, designed, executed and analysed all experiments. AY performed the simulations and analysed the data. GL performed the embryo collection. AL wrote the computational method sarlacc with the help of CTL and FB. MI and DG provided the hiPSCs. CEL helped conceive this study, and helped to design and optimize the CELLO-seq protocol. JM conceived and supervised this study. JM and RVB wrote the final version of the manuscript. All authors commented on the final manuscript.

## Acknowledgements

We thank the Sanger cytometry core facility for sorting of the hiPSCs. We thank the Sanger and CRUK sequencing facility for sequencing the NGS data. We thank Vasavi Sundaram for fruitful discussion. We thank Poppy Gould, Wolf Reik and Donal O’Carroll for helpful comments on the manuscript.

