## Supplemental Files for "Transposable element expression at unique loci in single cells with CELLO-seq"

#### Supplemental Figures

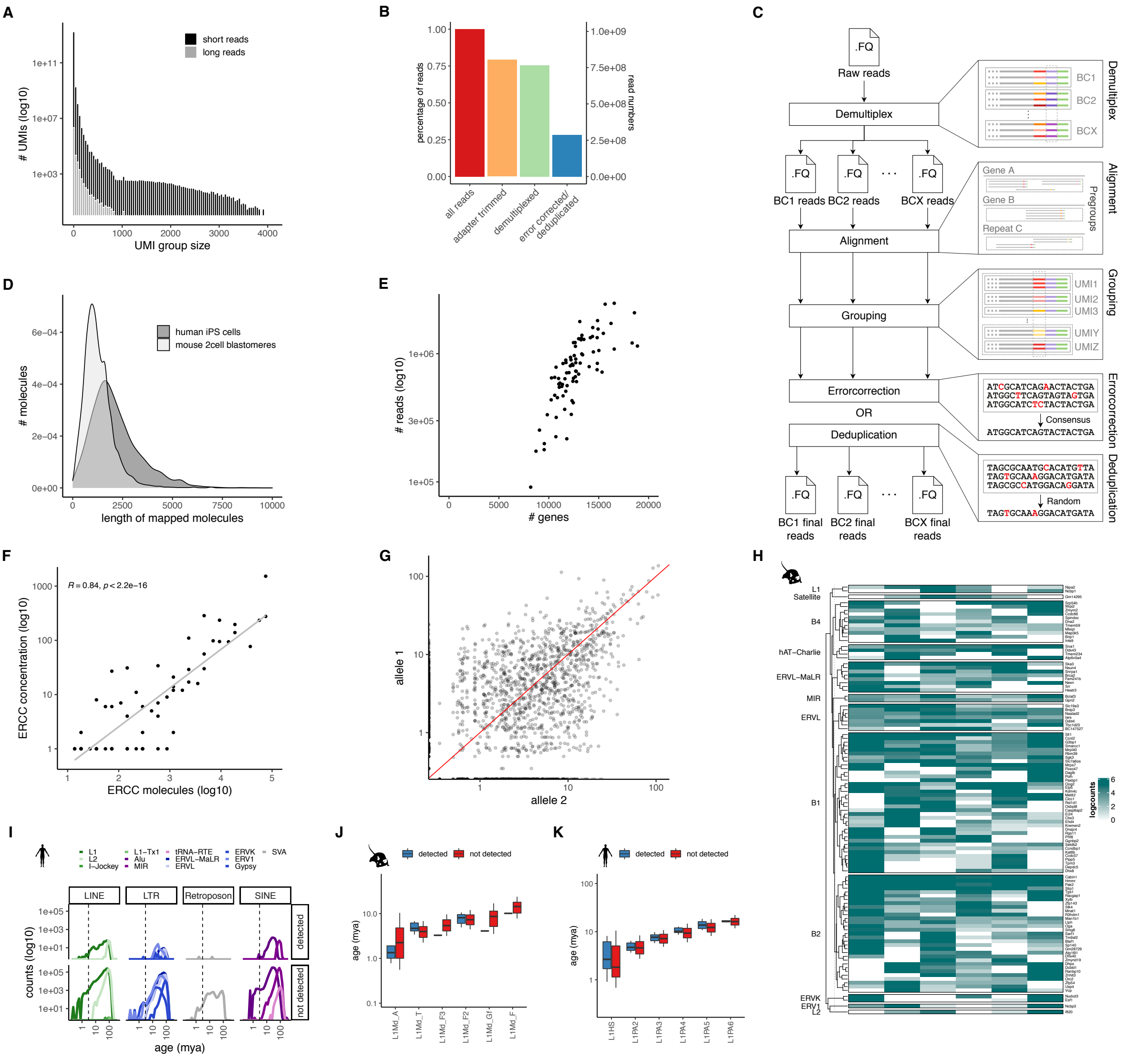


*Fig S1. **CELLO-seq library properties and isoform analysis.** (A) histogram of number of UMIs (log10) (y-axis) by UMI group size (x-axis) for mouse 2-cell blastomere CELLO-seq data either sequenced on a MinION flow cell (long reads, light grey) or Illumina Hiseq2500 (short reads, black). (B) bargraph of the percentage (y-axis left) read numbers (y-axis right) of total reads = red, after adapter trimming = orange, demultiplexing = light green as well as error correction or deduplication = light blue for mouse 2-cell embryo dataset. (C) schematic of sarlacc workflow. We demultiplexed the samples based on the sample barcodes by grouping barcodes that have a Levenshtein distance below the grouping threshold. To reduce the search space for the UMI grouping, we performed pre-grouping by mapping the reads to the relevant transcriptome. We grouped the reads by their UMI sequence and either error corrected the reads by taking the consensus sequence from multiple sequence alignment of up to 50 reads in the true UMI group, or by picking a random read from the UMI group in deduplication mode. For this study we used error corrected reads and aligned them back to the genome. (D) density plot of number of molecules (y-axis) by length of mapped molecules (x-axis) for hiPSCs dark grey) and mouse 2-cell blastomeres (light grey). (E) scatter plot of number of reads (log10) (y-axis) versus number of genes in 96 human iPSC short read sequencing data. These are from Smart-seq2 libraries sequenced by Illumina*

*HiSeq4000 at 150bp PE. (F) Pearson correlation of ERCC concentration (log10) (y-axis) from with ERCC molecules (log10) (x-axis) from mouse 2-cell data. Correlation coefficient and p-value shown. (G) scatter plot of read logcount correlation between allele1 (y-axis) and allele2 (x-axis) of CELLO-seq hiPSCs calculated using all heterozygous genic SNPs. Red line intercept=0 and slope=1. (H) Heatmap of logcounts of all TE-derived isoforms in mouse 2-cell embryos, with rows clustered by TE family and gene names shown as rownames. (I) frequency plot of number of repeats (x-axis) by age of TE (mya) (y-axis). Upper panel shows TEs we mapped in our study to hiPSCs and lower panel shows all repeats from mouse UCSC repeatmasker annotation (Smit et al., 2013-2015). TEs are grouped by TE class and colored by TE family. (J) boxplots of age (mya) of TEs (y-axis) of young L1Md elements which were either detected (red) in our 2-cell mouse dataset or not (dark blue). (K) boxplots of age (mya) of TEs (y-axis) of young L1HS and L1PA elements which were either detected (red) in our hiPSC dataset or not (dark blue). The boxplots shown in J, K show the median, first and third quartiles as a box, and the whiskers indicate the most extreme data point within 1.5 lengths of the box.*

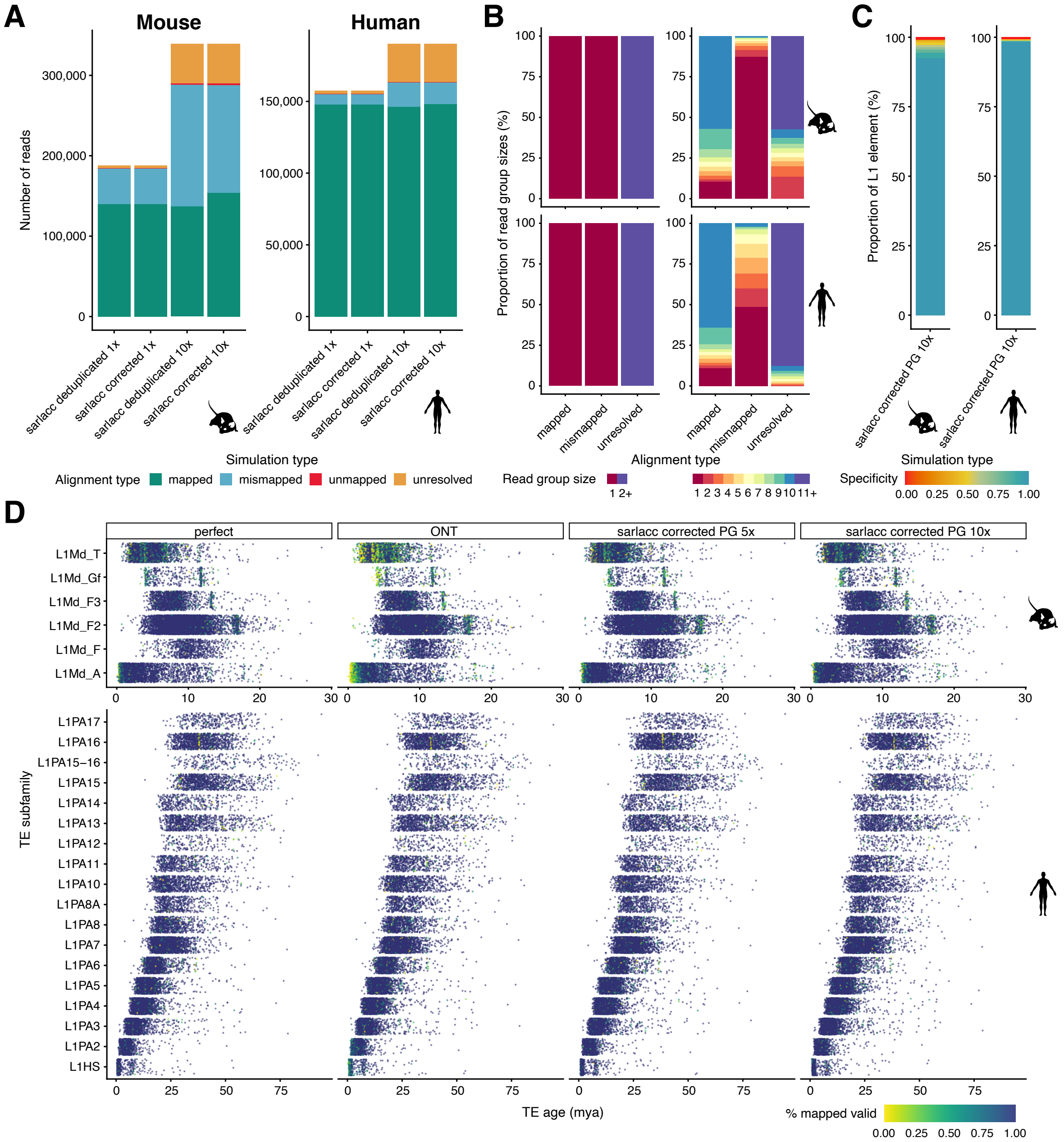


***Figure S2 Simulation of correct mapping of young L1 in mouse and human genome.*** *(A) bargraph showing the number of reads (y-axis) by the simulation type with either 1x or 10x coverage (x-axis), color-coded by alignment type with mapped = turquoise, read at correct location after mapping with minimap2 to the genome, mismapped = light blue, read maps at wrong location, unmapped = red, read not mapped, unresolved = yellow, group has more than one molecule present and group cannot be resolved to a unique read. Left is showing mouse L1s, right is showing human L1s. (B) bargraph of proportion of read group sizes (y-axis) by alignment type (x-axis), left showing 1x read coverage, right showing 10x read coverage. Color-coded by group size with red = 1, orange= 2, light orange = 3, yellow = 4, light green = 5, blue = 6+. Upper panel showing mouse data, lower panel showing human data. (C) Stacked bargraph showing proportion of L1 elements (y-axis) by simulation type using 10x read coverage (x-axis), colored by specificity score with red being 0% specificity and light blue being 100% specificity for mouse L1 left panel and human L1 right panel. (D) Jitter plot of TE subfamily (y-axis) by TE age (million years ago) grouped by simulation type and colored by % of mapped reads with yellow being 0% mapped and dark blue being 100% mapped. Mouse L1 top panel and human L1 bottom panel. Simulation type: perfect = perfect read identity, ONT = ONT read*

*identity, ONT 5x = ONT read identity with 5x coverage each read, sarlacc corrected 5x = ONT read identity score, 5x coverage run through sarlacc using error correction, sarlacc corrected10x = ONT read identity score, 10x coverage run through sarlacc using error correction, sarlacc deduplicated 5x = ONT read identity score, 5x coverage run through sarlacc using deduplication by randomly choosing 1 read. PG= perfect grouping. (D) Jitter plot of TE subfamily (y-axis) by TE age (million years ago) grouped by simulation type and colored by % of mapped reads with yellow being 0% mapped and dark blue being 100% mapped. Mouse L1 top panel and human L1 bottom panel.*

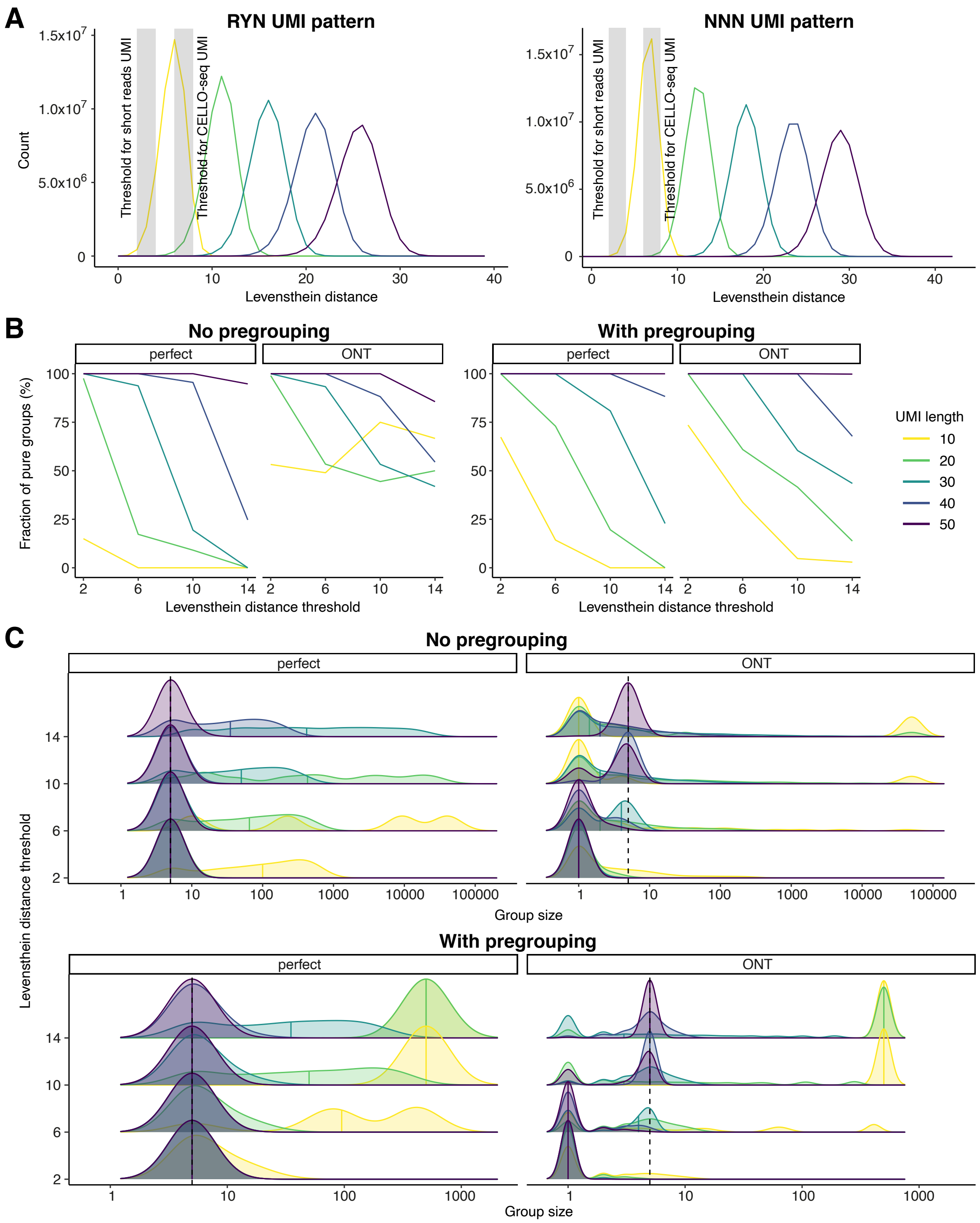


***Fig S3 UMI simulations.*** *(A) Distribution of evaluation of Levenshtein distance (x-axis) based on UMI length with RYN pattern (left) or NNN pattern (right). Light grey bar shows distance threshold for grouping of reads by UMIs used for most short read UMIs or CELLO-seq. (B) Line graph of fraction of pure groups (y-axis) by levenshtein distance (x-axis) by UMI group, either with perfect read identity or ONT read identity. On the left is the line graph of UMI simulations without any pregrouping by mapping. On the right the line graph is UMI simulation where pregrouping was performed by random assignment of true UMI sequences into groups of 100 unique UMIs. (C) distribution plot of UMI group sizes (x-axis) by levenshtein distance threshold (y-axis) based on UMI length, with perfect of ONT read identity and no pregrouping (left) or pregrouping (right).*

*UMI length 10 = yellow, UMI length 20 = green, UMI length 30 = light blue, UMI length 40 = dark blue, UMI length 50 = dark purple.*

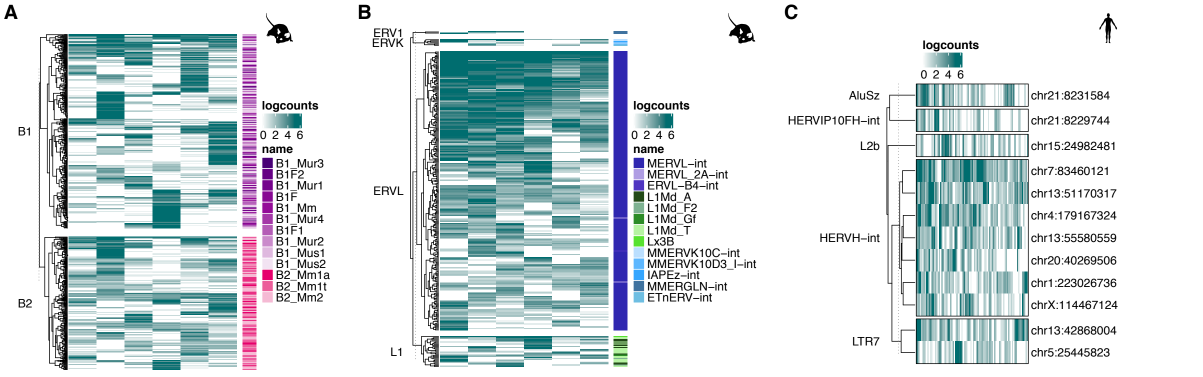


***Fig S4. CELLO-seq to study functional correlation of TEs and gene expression.*** *(A) Heatmap of logcounts of all SINE B1 and B2 elements in mouse 2-cell embryos, with rows clustered by SINE family and color coded by TE subfamily: pink = SINE B2, purple = SINE B1. (B) Heatmap of logcounts of full-length (>5000nt) elements in mouse 2-cell embryos, with rows clustered by TE family and color coded by TE subfamily: dark purple = ERVL, light blue = ERVK, L1 = green, dark blue = ERV1. (C) Heatmap of logcounts of highest expressed (mean expression over all cells > 1) elements in hiPSCs with rows clustered by TE subfamily and rownames of unique TE loci.*

***Table S1. Specificity score of L1 elements used in this study.*** *We use only L1 element with specificity score of >80%.*

| | strand | jc_distance | mya | length | specificity |
|---|---|---|---|---|---|
| **human L1** | | | | | |
| L1PA2_chr3_24089455-24095487 | - | 0.014 | 3.212 | 6032 | 1.00 |
| L1PA6_chr3_142135407-142141502 | - | 0.068 | 15.452 | 6095 | 1.00 |
| L1PA3_chr4_61743722-61749869 | + | 0.045 | 10.305 | 6147 | 1.00 |
| L1PA7_chr4_76477676-76484153 | + | 0.090 | 20.503 | 6477 | 1.00 |
| L1HS_chr6_93865739-93871783 | - | 0.008 | 1.828 | 6044 | 1.00 |
| L1PA2_chr6_127593293-127599292 | + | 0.026 | 6.014 | 5999 | 1.00 |
| L1HS_chr6_156324981-156331010 | - | 0.009 | 2.058 | 6029 | 1.00 |
| L1PA2_chr7_30636809-30642830 | - | 0.019 | 4.374 | 6021 | 1.00 |
| L1PA12_chrX_98910031-98916276 | - | 0.188 | 42.643 | 6245 | 1.00 |
| L1PA2_chr11_43346102-43352118 | - | 0.013 | 2.980 | 6016 | 1.00 |
| L1PA7_chr11_85542992-85549441 | - | 0.127 | 28.910 | 6449 | 1.00 |
| L1HS_chr18_72966527-72972556 | - | 0.009 | 2.058 | 6029 | 1.00 |
| L1HS_chr20_11632780-11638837 | + | 0.004 | 0.912 | 6057 | 1.00 |
| L1PA8_chr20_39969159-39975570 | - | 0.127 | 28.910 | 6411 | 1.00 |
| **mouse L1** | | | | | |
| L1Md_T_chr1_17383284-17390159 | - | 0.031 | 3.402 | 6875 | 1.00 |
| L1Md_T_chr2_14118838-14125461 | - | 0.055 | 6.107 | 6623 | 1.00 |
| L1Md_A_chr2_63718187-63725386 | - | 0.008 | 0.894 | 7199 | 1.00 |
| L1Md_T_chr2_136863719-136870367 | - | 0.058 | 6.467 | 6648 | 1.00 |
| L1Md_A_chr3_78332502-78339863 | - | 0.015 | 1.684 | 7361 | 0.83 |
| L1Md_T_chr3_94146299-94152497 | - | 0.045 | 5.038 | 6198 | 1.00 |
| L1Md_T_chr3_148704254-148710467 | - | 0.038 | 4.216 | 6213 | 1.00 |
| L1Md_A_chr3_158730068-158739645 | - | 0.038 | 4.216 | 9577 | 1.00 |
| L1Md_A_chr4_9585467-9591708 | + | 0.017 | 1.911 | 6241 | 1.00 |
| L1Md_A_chr7_5269032-5276386 | - | 0.009 | 1.006 | 7354 | 0.80 |
| L1Md_T_chr9_12739974-12745831 | + | 0.036 | 3.983 | 5857 | 1.00 |
| L1Md_T_chr9_93013303-93021296 | - | 0.061 | 6.828 | 7993 | 1.00 |
| L1Md_A_chr10_27812548-27819961 | - | 0.015 | 1.684 | 7413 | 1.00 |
| L1Md_F2_chr10_50808787-50814870 | - | 0.075 | 8.288 | 6083 | 1.00 |
| L1Md_T_chr12_55632616-55638687 | + | 0.035 | 3.866 | 6071 | 1.00 |
| L1Md_T_chr13_9832021-9838665 | - | 0.041 | 4.567 | 6644 | 1.00 |
| L1Md_A_chr13_64582655-64589456 | - | 0.013 | 1.457 | 6801 | 1.00 |
| L1Md_T_chr13_64642649-64649318 | - | 0.047 | 5.275 | 6669 | 1.00 |
| L1Md_T_chr17_69323423-69330344 | + | 0.037 | 4.099 | 6921 | 1.00 |
| L1Md_T_chr18_73681328-73687549 | - | 0.030 | 3.286 | 6221 | 1.00 |
| L1Md_A_chr18_89136300-89142340 | + | 0.015 | 1.684 | 6040 | 1.00 |

### Table S2. Oligonucleotide sequences used in this manuscript

| Name | Sequence (5'-3') |
|---|---|
| Illumina_i7_N723 | CAAGCAGAAGACGGCATACGAGATGAGCGCTAGTCTCGTGGGCTCGG |
| Illumina_i5_S520 | AATGATACGGCGACCACCGAGATCTACACAAGGCTATTCGTCGGCAGCGTC |
| Tn5_ME_RC | Phos-CTGTCTCTTATACACATCT |
| NexteR1 | CGTCGGCAGCGTCAGATGTGTATAAGAGACAG |
| NexteR2 | GTCTCGTGGGCTCGGAGATGTGTATAAGAGACAG |
| oligo_dT | Phos-TGGCGTAGCGGGTTCGAGCGCACCGCAGGGTATCCGGCTATTTTTTTTTTTTTTTTTTTTTTTTTTTVN |
| TSO_RNA_Amine | TCGTCGGCAGCGTCAGATGTGTATAAGAGACArGrGrGAmn |
| splint_top | TAGCCGGATACCCTGCGGTGCGCTCGAACCCGCTACGCCATGTAAAACGACGGCCAGTInvdT |
| splintlig_spc_idx1 | GTCTCGTGGGCTCGGAGATGTGTATAAGAGACAGAAGAAAGTTGTCGGTGTCTTTGAGTGGTATCNRYNRYNRYNRYNRYNRYNRYNACTGGCCGTCGTTTTACATGGCGTAGCGGGTTCGAGCGCACCGCAGGGTATCCGGCTATTTTTTTTTTTTTTT |
| splintlig_spc_idx2 | GTCTCGTGGGCTCGGAGATGTGTATAAGAGACAGTCGATTCCGTTTGTAGTCGTCTGTAGTGGTATCNRYNRYNRYNRYNRYNRYNRYNACTGGCCGTCGTTTTACATGGCGTAGCGGGTTCGAGCGCACCGCAGGGTATCCGGCTATTTTTTTTTTTTTTT |
| splintlig_spc_idx3 | GTCTCGTGGGCTCGGAGATGTGTATAAGAGACAGGAGTCTTGTGTCCCAGTTACCAGGAGTGGTATCNRYNRYNRYNRYNRYNRYNRYNACTGGCCGTCGTTTTACATGGCGTAGCGGGTTCGAGCGCACCGCAGGGTATCCGGCTATTTTTTTTTTTTTTT |
| splintlig_spc_idx4 | GTCTCGTGGGCTCGGAGATGTGTATAAGAGACAGTTCGGATTCTATCGTGTTTCCCTAAGTGGTATCNRYNRYNRYNRYNRYNRYNRYNACTGGCCGTCGTTTTACATGGCGTAGCGGGTTCGAGCGCACCGCAGGGTATCCGGCTATTTTTTTTTTTTTTT |
| splintlig_spc_idx5 | GTCTCGTGGGCTCGGAGATGTGTATAAGAGACAGCTTGTCCAGGGTTTGTGTAACCTTAGTGGTATCNRYNRYNRYNRYNRYNRYNRYNACTGGCCGTCGTTTTACATGGCGTAGCGGGTTCGAGCGCACCGCAGGGTATCCGGCTATTTTTTTTTTTTTTT |
| splintlig_spc_idx6 | GTCTCGTGGGCTCGGAGATGTGTATAAGAGACAGTTCTCGCAAAGGCAGAAAGTAGTCAGTGGTATCNRYNRYNRYNRYNRYNRYNRYNACTGGCCGTCGTTTTACATGGCGTAGCGGGTTCGAGCGCACCGCAGGGTATCCGGCTATTTTTTTTTTTTTTT |
| splintlig_spc_idx7 | GTCTCGTGGGCTCGGAGATGTGTATAAGAGACAGGTGTTACCGTGGGAATGAATCCTTAGTGGTATCNRYNRYNRYNRYNRYNRYNRYNACTGGCCGTCGTTTTACATGGCGTAGCGGGTTCGAGCGCACCGCAGGGTATCCGGCTATTTTTTTTTTTTTTT |
| splintlig_spc_idx8 | GTCTCGTGGGCTCGGAGATGTGTATAAGAGACAGTTCAGGGAACAAACCAAGTTACGTAGTGGTATCNRYNRYNRYNRYNRYNRYNRYNACTGGCCGTCGTTTTACATGGCGTAGCGGGTTCGAGCGCACCGCAGGGTATCCGGCTATTTTTTTTTTTTTTT |
| splintlig_spc_idx9 | GTCTCGTGGGCTCGGAGATGTGTATAAGAGACAGAACTAGGCACAGCGAGTCTTGGTTAGTGGTATCNRYNRYNRYNRYNRYNRYNRYNACTGGCCGTCGTTTTACATGGCGTAGCGGGTTCGAGCGCACCGCAGGGTATCCGGCTATTTTTTTTTTTTTTT |
| splintlig_spc_idx10 | GTCTCGTGGGCTCGGAGATGTGTATAAGAGACAGAAGCGTTGAAACCTTTGTCCTCTCAGTGGTATCNRYNRYNRYNRYNRYNRYNRYNACTGGCCGTCGTTTTACATGGCGTAGCGGGTTCGAGCGCACCGCAGGGTATCCGGCTATTTTTTTTTTTTTTT |
| splintlig_spc_idx11 | GTCTCGTGGGCTCGGAGATGTGTATAAGAGACAGGTTTCATCTATCGGAGGGAATGGAAGTGGTATCNRYNRYNRYNRYNRYNRYNRYNACTGGCCGTCGTTTTACATGGCGTAGCGGGTTCGAGCGCACCGCAGGGTATCCGGCTATTTTTTTTTTTTTTT |
| splintlig_spc_idx12 | GTCTCGTGGGCTCGGAGATGTGTATAAGAGACAGCAGGTAGAAAGAAGCAGAATCGGAAGTGGTATCNRYNRYNRYNRYNRYNRYNRYNACTGGCCGTCGTTTTACATGGCGTAGCGGGTTCGAGCGCACCGCAGGGTATCCGGCTATTTTTTTTTTTTTTT |
| splint_5pr_lig_idx13 | GTCTCGTGGGCTCGGAGATGTGTATAAGAGACAGAACGAGTCTCTTGGGACCCATAGAAGTGGTATCNRYNRYNRYNRYNRYNRYNRYNACTGGCCGTCGTTTTACATGGCGTAGCGGGTTCGAGCGCACCGCAGGGTATCCGGCTATTTTTTTTTTTTTTT |
| splint_5pr_lig_idx14 | GTCTCGTGGGCTCGGAGATGTGTATAAGAGACAGAGAACGACTTCCATACTCGTGTGAAGTGGTATCNRYNRYNRYNRYNRYNRYNRYNACTGGCCGTCGTTTTACATGGCGTAGCGGGTTCGAGCGCACCGCAGGGTATCCGGCTATTTTTTTTTTTTTTT |
| splint_5pr_lig_idx15 | GTCTCGTGGGCTCGGAGATGTGTATAAGAGACAGAGGTCTACCTCGCTAACACCACTGAGTGGTATCNRYNRYNRYNRYNRYNRYNRYNACTGGCCGTCGTTTTACATGGCGTAGCGGGTTCGAGCGCACCGCAGGGTATCCGGCTATTTTTTTTTTTTTTT |
| splint_5pr_lig_idx16 | GTCTCGTGGGCTCGGAGATGTGTATAAGAGACAGCGTCAACTGACAGTGGTTCGTACTAGTGGTATCNRYNRYNRYNRYNRYNRYNRYNACTGGCCGTCGTTTTACATGGCGTAGCGGGTTCGAGCGCACCGCAGGGTATCCGGCTATTTTTTTTTTTTTTT |
| splint_5pr_lig_idx17 | GTCTCGTGGGCTCGGAGATGTGTATAAGAGACAGACCCTCCAGGAAAGTACCTCTGATAGTGGTATCNRYNRYNRYNRYNRYNRYNRYNACTGGCCGTCGTTTTACATGGCGTAGCGGGTTCGAGCGCACCGCAGGGTATCCGGCTATTTTTTTTTTTTTTT |
| splint_5pr_lig_idx18 | GTCTCGTGGGCTCGGAGATGTGTATAAGAGACAGCCAAACCCAACAACCTAGATAGGCAGTGGTATCNRYNRYNRYNRYNRYNRYNRYNACTGGCCGTCGTTTTACATGGCGTAGCGGGTTCGAGCGCACCGCAGGGTATCCGGCTATTTTTTTTTTTTTTT |
| splint_5pr_lig_idx19 | GTCTCGTGGGCTCGGAGATGTGTATAAGAGACAGGTTCCTCGTGCAGTGTCAAGAGATAGTGGTATCNRYNRYNRYNRYNRYNRYNRYNACTGGCCGTCGTTTTACATGGCGTAGCGGGTTCGAGCGCACCGCAGGGTATCCGGCTATTTTTTTTTTTTTTT |
| splint_5pr_lig_idx20 | GTCTCGTGGGCTCGGAGATGTGTATAAGAGACAGGAGCCTCTCATTGTCCGTTCTCTAAGTGGTATCNRYNRYNRYNRYNRYNRYNRYNACTGGCCGTCGTTTTACATGGCGTAGCGGGTTCGAGCGCACCGCAGGGTATCCGGCTATTTTTTTTTTTTTTT |
| splint_5pr_lig_idx21 | GTCTCGTGGGCTCGGAGATGTGTATAAGAGACAGTTGCGTCCTGTTACGAGAACTCATAGTGGTATCNRYNRYNRYNRYNRYNRYNRYNACTGGCCGTCGTTTTACATGGCGTAGCGGGTTCGAGCGCACCGCAGGGTATCCGGCTATTTTTTTTTTTTTTT |
| splint_5pr_lig_idx22 | GTCTCGTGGGCTCGGAGATGTGTATAAGAGACAGACCACTGCCATGTATCAAAGTACGAGTGGTATCNRYNRYNRYNRYNRYNRYNRYNACTGGCCGTCGTTTTACATGGCGTAGCGGGTTCGAGCGCACCGCAGGGTATCCGGCTATTTTTTTTTTTTTTT |
| splint_5pr_lig_idx23 | GTCTCGTGGGCTCGGAGATGTGTATAAGAGACAGCTTACTACCCAGTGAACCTCCTCGAGTGGTATCNRYNRYNRYNRYNRYNRYNRYNACTGGCCGTCGTTTTACATGGCGTAGCGGGTTCGAGCGCACCGCAGGGTATCCGGCTATTTTTTTTTTTTTTT |
| splint_5pr_lig_idx24 | GTCTCGTGGGCTCGGAGATGTGTATAAGAGACAGGCATAGTTCTGCATGATGGGTTAGAGTGGTATCNRYNRYNRYNRYNRYNRYNRYNACTGGCCGTCGTTTTACATGGCGTAGCGGGTTCGAGCGCACCGCAGGGTATCCGGCTATTTTTTTTTTTTTTT |
| splint_5pr_lig_idx25 | GTCTCGTGGGCTCGGAGATGTGTATAAGAGACAGGTAAGTTGGGTATGCAACGCAATGAGTGGTATCNRYNRYNRYNRYNRYNRYNRYNACTGGCCGTCGTTTTACATGGCGTAGCGGGTTCGAGCGCACCGCAGGGTATCCGGCTATTTTTTTTTTTTTTT |
| splint_5pr_lig_idx26 | GTCTCGTGGGCTCGGAGATGTGTATAAGAGACAGCATACAGCGACTACGCATTCTCATAGTGGTATCNRYNRYNRYNRYNRYNRYNRYNACTGGCCGTCGTTTTACATGGCGTAGCGGGTTCGAGCGCACCGCAGGGTATCCGGCTATTTTTTTTTTTTTTT |
| splint_5pr_lig_idx27 | GTCTCGTGGGCTCGGAGATGTGTATAAGAGACAGCGACGGTTAGATTCACCTCTTACAAGTGGTATCNRYNRYNRYNRYNRYNRYNRYNACTGGCCGTCGTTTTACATGGCGTAGCGGGTTCGAGCGCACCGCAGGGTATCCGGCTATTTTTTTTTTTTTTT |
| splint_5pr_lig_idx28 | GTCTCGTGGGCTCGGAGATGTGTATAAGAGACAGTGAAACCTAAGAAGGCACCGTATCAGTGGTATCNRYNRYNRYNRYNRYNRYNRYNACTGGCCGTCGTTTTACATGGCGTAGCGGGTTCGAGCGCACCGCAGGGTATCCGGCTATTTTTTTTTTTTTTT |
| splint_5pr_lig_idx29 | GTCTCGTGGGCTCGGAGATGTGTATAAGAGACAGCTAGACACCTTGGGTTGACAGACCAGTGGTATCNRYNRYNRYNRYNRYNRYNRYNACTGGCCGTCGTTTTACATGGCGTAGCGGGTTCGAGCGCACCGCAGGGTATCCGGCTATTTTTTTTTTTTTTT |
| splint_5pr_lig_idx30 | GTCTCGTGGGCTCGGAGATGTGTATAAGAGACAGTCAGTGAGGATCTACTTCGACCCAAGTGGTATCNRYNRYNRYNRYNRYNRYNRYNACTGGCCGTCGTTTTACATGGCGTAGCGGGTTCGAGCGCACCGCAGGGTATCCGGCTATTTTTTTTTTTTTTT |
| splint_5pr_lig_idx31 | GTCTCGTGGGCTCGGAGATGTGTATAAGAGACAGTGCGTACAGCAATCAGTTACATTGAGTGGTATCNRYNRYNRYNRYNRYNRYNRYNACTGGCCGTCGTTTTACATGGCGTAGCGGGTTCGAGCGCACCGCAGGGTATCCGGCTATTTTTTTTTTTTTTT |
| splint_5pr_lig_idx32 | GTCTCGTGGGCTCGGAGATGTGTATAAGAGACAGCCAGTAGAAGTCCGACAACGTCATAGTGGTATCNRYNRYNRYNRYNRYNRYNRYNACTGGCCGTCGTTTTACATGGCGTAGCGGGTTCGAGCGCACCGCAGGGTATCCGGCTATTTTTTTTTTTTTTT |
| splint_5pr_lig_idx33 | GTCTCGTGGGCTCGGAGATGTGTATAAGAGACAGCAGACTTGGTACGGTTGGGTAACTAGTGGTATCNRYNRYNRYNRYNRYNRYNRYNACTGGCCGTCGTTTTACATGGCGTAGCGGGTTCGAGCGCACCGCAGGGTATCCGGCTATTTTTTTTTTTTTTT |
| splint_5pr_lig_idx34 | GTCTCGTGGGCTCGGAGATGTGTATAAGAGACAGGGACGAAGAACTCAAGTCAAAGGCAGTGGTATCNRYNRYNRYNRYNRYNRYNRYNACTGGCCGTCGTTTTACATGGCGTAGCGGGTTCGAGCGCACCGCAGGGTATCCGGCTATTTTTTTTTTTTTTT |
| splint_5pr_lig_idx35 | GTCTCGTGGGCTCGGAGATGTGTATAAGAGACAGCTACTTACGAAGCTGAGGGACTGCAGTGGTATCNRYNRYNRYNRYNRYNRYNRYNACTGGCCGTCGTTTTACATGGCGTAGCGGGTTCGAGCGCACCGCAGGGTATCCGGCTATTTTTTTTTTTTTTT |
| splint_5pr_lig_idx36 | GTCTCGTGGGCTCGGAGATGTGTATAAGAGACAGATGTCCCAGTTAGAGGAGGAAACAAGTGGTATCNRYNRYNRYNRYNRYNRYNRYNACTGGCCGTCGTTTTACATGGCGTAGCGGGTTCGAGCGCACCGCAGGGTATCCGGCTATTTTTTTTTTTTTTT |
| splint_5pr_lig_idx37 | GTCTCGTGGGCTCGGAGATGTGTATAAGAGACAGGCTTGCGATTGATGCTTAGTATCAAGTGGTATCNRYNRYNRYNRYNRYNRYNRYNACTGGCCGTCGTTTTACATGGCGTAGCGGGTTCGAGCGCACCGCAGGGTATCCGGCTATTTTTTTTTTTTTTT |
| splint_5pr_lig_idx38 | GTCTCGTGGGCTCGGAGATGTGTATAAGAGACAGACCACAGGAGGACGATACAGAGAAAGTGGTATCNRYNRYNRYNRYNRYNRYNRYNACTGGCCGTCGTTTTACATGGCGTAGCGGGTTCGAGCGCACCGCAGGGTATCCGGCTATTTTTTTTTTTTTTT |
| splint_5pr_lig_idx39 | GTCTCGTGGGCTCGGAGATGTGTATAAGAGACAGCCACAGTGTCAACTAGAGCCTCTCAGTGGTATCNRYNRYNRYNRYNRYNRYNRYNACTGGCCGTCGTTTTACATGGCGTAGCGGGTTCGAGCGCACCGCAGGGTATCCGGCTATTTTTTTTTTTTTTT |
| splint_5pr_lig_idx40 | GTCTCGTGGGCTCGGAGATGTGTATAAGAGACAGTAGTTTGGATGACCAAGGATAGCCAGTGGTATCNRYNRYNRYNRYNRYNRYNRYNACTGGCCGTCGTTTTACATGGCGTAGCGGGTTCGAGCGCACCGCAGGGTATCCGGCTATTTTTTTTTTTTTTT |
| splint_5pr_lig_idx41 | GTCTCGTGGGCTCGGAGATGTGTATAAGAGACAGGGAGTTCGTCCAGAGAAGTACACGAGTGGTATCNRYNRYNRYNRYNRYNRYNRYNACTGGCCGTCGTTTTACATGGCGTAGCGGGTTCGAGCGCACCGCAGGGTATCCGGCTATTTTTTTTTTTTTTT |
| splint_5pr_lig_idx42 | GTCTCGTGGGCTCGGAGATGTGTATAAGAGACAGCTACGTGTAAGGCATACCTGCCAGAGTGGTATCNRYNRYNRYNRYNRYNRYNRYNACTGGCCGTCGTTTTACATGGCGTAGCGGGTTCGAGCGCACCGCAGGGTATCCGGCTATTTTTTTTTTTTTTT |
| splint_5pr_lig_idx43 | GTCTCGTGGGCTCGGAGATGTGTATAAGAGACAGCTTTCGTTGTTGACTCGACGGTAGAGTGGTATCNRYNRYNRYNRYNRYNRYNRYNACTGGCCGTCGTTTTACATGGCGTAGCGGGTTCGAGCGCACCGCAGGGTATCCGGCTATTTTTTTTTTTTTTT |
| splint_5pr_lig_idx44 | GTCTCGTGGGCTCGGAGATGTGTATAAGAGACAGAGTAGAAAGGGTTCCTTCCCACTCAGTGGTATCNRYNRYNRYNRYNRYNRYNRYNACTGGCCGTCGTTTTACATGGCGTAGCGGGTTCGAGCGCACCGCAGGGTATCCGGCTATTTTTTTTTTTTTTT |
| splint_5pr_lig_idx45 | GTCTCGTGGGCTCGGAGATGTGTATAAGAGACAGGATCCAACAGAGATGCCTTCAGTGAGTGGTATCNRYNRYNRYNRYNRYNRYNRYNACTGGCCGTCGTTTTACATGGCGTAGCGGGTTCGAGCGCACCGCAGGGTATCCGGCTATTTTTTTTTTTTTTT |
| splint_5pr_lig_idx46 | GTCTCGTGGGCTCGGAGATGTGTATAAGAGACAGGCTGTGTTCCACTTCATTCTCCTGAGTGGTATCNRYNRYNRYNRYNRYNRYNRYNACTGGCCGTCGTTTTACATGGCGTAGCGGGTTCGAGCGCACCGCAGGGTATCCGGCTATTTTTTTTTTTTTTT |
| splint_5pr_lig_idx47 | GTCTCGTGGGCTCGGAGATGTGTATAAGAGACAGGTGCAACTTTCCCACAGGTAGTTCAGTGGTATCNRYNRYNRYNRYNRYNRYNRYNACTGGCCGTCGTTTTACATGGCGTAGCGGGTTCGAGCGCACCGCAGGGTATCCGGCTATTTTTTTTTTTTTTT |
| splint_5pr_lig_idx48 | GTCTCGTGGGCTCGGAGATGTGTATAAGAGACAGCATCTGGAACGTGGTACACCTGTAAGTGGTATCNRYNRYNRYNRYNRYNRYNRYNACTGGCCGTCGTTTTACATGGCGTAGCGGGTTCGAGCGCACCGCAGGGTATCCGGCTATTTTTTTTTTTTTTT |
| splint_5pr_lig_idx49 | GTCTCGTGGGCTCGGAGATGTGTATAAGAGACAGACTGGTGCAGCTTTGAACATCTAGAGTGGTATCNRYNRYNRYNRYNRYNRYNRYNACTGGCCGTCGTTTTACATGGCGTAGCGGGTTCGAGCGCACCGCAGGGTATCCGGCTATTTTTTTTTTTTTTT |
| splint_5pr_lig_idx50 | GTCTCGTGGGCTCGGAGATGTGTATAAGAGACAGATGGACTTTGGTAACTTCCTGCGTAGTGGTATCNRYNRYNRYNRYNRYNRYNRYNACTGGCCGTCGTTTTACATGGCGTAGCGGGTTCGAGCGCACCGCAGGGTATCCGGCTATTTTTTTTTTTTTTT |
| splint_5pr_lig_idx51 | GTCTCGTGGGCTCGGAGATGTGTATAAGAGACAGGTTGAATGAGCCTACTGGGTCCTCAGTGGTATCNRYNRYNRYNRYNRYNRYNRYNACTGGCCGTCGTTTTACATGGCGTAGCGGGTTCGAGCGCACCGCAGGGTATCCGGCTATTTTTTTTTTTTTTT |
| splint_5pr_lig_idx52 | GTCTCGTGGGCTCGGAGATGTGTATAAGAGACAGTGAGAGACAAGATTGTTCGTGGACAGTGGTATCNRYNRYNRYNRYNRYNRYNRYNACTGGCCGTCGTTTTACATGGCGTAGCGGGTTCGAGCGCACCGCAGGGTATCCGGCTATTTTTTTTTTTTTTT |
| splint_5pr_lig_idx53 | GTCTCGTGGGCTCGGAGATGTGTATAAGAGACAGAGATTCAGACCGTCTCATGCAAAGCAGTGGTATCNRYNRYNRYNRYNRYNRYNRYNACTGGCCGTCGTTTTACATGGCGTAGCGGGTTCGAGCGCACCGCAGGGTATCCGGCTATTTTTTTTTTTTTTT |
| splint_5pr_lig_idx54 | GTCTCGTGGGCTCGGAGATGTGTATAAGAGACAGCAAGAGCTTTGACTAAGGAGCATGAGTGGTATCNRYNRYNRYNRYNRYNRYNRYNACTGGCCGTCGTTTTACATGGCGTAGCGGGTTCGAGCGCACCGCAGGGTATCCGGCTATTTTTTTTTTTTTTT |
| splint_5pr_lig_idx55 | GTCTCGTGGGCTCGGAGATGTGTATAAGAGACAGTGGAAGATGAGACCCTGATCTACGAGTGGTATCNRYNRYNRYNRYNRYNRYNRYNACTGGCCGTCGTTTTACATGGCGTAGCGGGTTCGAGCGCACCGCAGGGTATCCGGCTATTTTTTTTTTTTTTT |
| splint_5pr_lig_idx56 | GTCTCGTGGGCTCGGAGATGTGTATAAGAGACAGTCACTACTCAACAGGTGGCATGAAAGTGGTATCNRYNRYNRYNRYNRYNRYNRYNACTGGCCGTCGTTTTACATGGCGTAGCGGGTTCGAGCGCACCGCAGGGTATCCGGCTATTTTTTTTTTTTTTT |
| splint_5pr_lig_idx57 | GTCTCGTGGGCTCGGAGATGTGTATAAGAGACAGGCTAGGTCAATCTCCTTCGGAAGTAGTGGTATCNRYNRYNRYNRYNRYNRYNRYNACTGGCCGTCGTTTTACATGGCGTAGCGGGTTCGAGCGCACCGCAGGGTATCCGGCTATTTTTTTTTTTTTTT |
| splint_5pr_lig_idx58 | GTCTCGTGGGCTCGGAGATGTGTATAAGAGACAGCAGGTTACTCCTCCGTGAGTCTGAAGTGGTATCNRYNRYNRYNRYNRYNRYNRYNACTGGCCGTCGTTTTACATGGCGTAGCGGGTTCGAGCGCACCGCAGGGTATCCGGCTATTTTTTTTTTTTTTT |
| splint_5pr_lig_idx59 | GTCTCGTGGGCTCGGAGATGTGTATAAGAGACAGTCAATCAAGAAGGGAAAGCAAGGTAGTGGTATCNRYNRYNRYNRYNRYNRYNRYNACTGGCCGTCGTTTTACATGGCGTAGCGGGTTCGAGCGCACCGCAGGGTATCCGGCTATTTTTTTTTTTTTTT |
| splint_5pr_lig_idx60 | GTCTCGTGGGCTCGGAGATGTGTATAAGAGACAGCATGTTCAACCAAGGCTTCTATGGAGTGGTATCNRYNRYNRYNRYNRYNRYNRYNACTGGCCGTCGTTTTACATGGCGTAGCGGGTTCGAGCGCACCGCAGGGTATCCGGCTATTTTTTTTTTTTTTT |
| splint_5pr_lig_idx61 | GTCTCGTGGGCTCGGAGATGTGTATAAGAGACAGAGAGGGTACTATGTGCCTCAACGCAGTGGTATCNRYNRYNRYNRYNRYNRYNRYNACTGGCCGTCGTTTTACATGGCGTAGCGGGTTCGAGCGCACCGCAGGGTATCCGGCTATTTTTTTTTTTTTTT |
| splint_5pr_lig_idx62 | GTCTCGTGGGCTCGGAGATGTGTATAAGAGACAGCACCCACACTTACTTCAGGACGTAAGTGGTATCNRYNRYNRYNRYNRYNRYNRYNACTGGCCGTCGTTTTACATGGCGTAGCGGGTTCGAGCGCACCGCAGGGTATCCGGCTATTTTTTTTTTTTTTT |
| splint_5pr_lig_idx63 | GTCTCGTGGGCTCGGAGATGTGTATAAGAGACAGTTCTGAAGTTCCTGGGTCTTGAACAGTGGTATCNRYNRYNRYNRYNRYNRYNRYNACTGGCCGTCGTTTTACATGGCGTAGCGGGTTCGAGCGCACCGCAGGGTATCCGGCTATTTTTTTTTTTTTTT |
| splint_5pr_lig_idx64 | GTCTCGTGGGCTCGGAGATGTGTATAAGAGACAGGACAGACACCGTTCATCGACTTTCAGTGGTATCNRYNRYNRYNRYNRYNRYNRYNACTGGCCGTCGTTTTACATGGCGTAGCGGGTTCGAGCGCACCGCAGGGTATCCGGCTATTTTTTTTTTTTTTT |
| splint_5pr_lig_idx65 | GTCTCGTGGGCTCGGAGATGTGTATAAGAGACAGTTCTCAGTCTTCCTCCAGACAAGGAGTGGTATCNRYNRYNRYNRYNRYNRYNRYNACTGGCCGTCGTTTTACATGGCGTAGCGGGTTCGAGCGCACCGCAGGGTATCCGGCTATTTTTTTTTTTTTTT |
| splint_5pr_lig_idx66 | GTCTCGTGGGCTCGGAGATGTGTATAAGAGACAGCCGATCCTTGTGGCTTCTAACTTCAGTGGTATCNRYNRYNRYNRYNRYNRYNRYNACTGGCCGTCGTTTTACATGGCGTAGCGGGTTCGAGCGCACCGCAGGGTATCCGGCTATTTTTTTTTTTTTTT |
| splint_5pr_lig_idx67 | GTCTCGTGGGCTCGGAGATGTGTATAAGAGACAGGTTTGTCATACTCGTGTGCTCACCAGTGGTATCNRYNRYNRYNRYNRYNRYNRYNACTGGCCGTCGTTTTACATGGCGTAGCGGGTTCGAGCGCACCGCAGGGTATCCGGCTATTTTTTTTTTTTTTT |
| splint_5pr_lig_idx68 | GTCTCGTGGGCTCGGAGATGTGTATAAGAGACAGGAATCTAAGCAAACACGAAGGTGGAGTGGTATCNRYNRYNRYNRYNRYNRYNRYNACTGGCCGTCGTTTTACATGGCGTAGCGGGTTCGAGCGCACCGCAGGGTATCCGGCTATTTTTTTTTTTTTTT |
| splint_5pr_lig_idx69 | GTCTCGTGGGCTCGGAGATGTGTATAAGAGACAGTACAGTCCGAGCCTCATGTGATCTAGTGGTATCNRYNRYNRYNRYNRYNRYNRYNACTGGCCGTCGTTTTACATGGCGTAGCGGGTTCGAGCGCACCGCAGGGTATCCGGCTATTTTTTTTTTTTTTT |
| splint_5pr_lig_idx70 | GTCTCGTGGGCTCGGAGATGTGTATAAGAGACAGACCGAGATCCTACGAATGGAGTGTAGTGGTATCNRYNRYNRYNRYNRYNRYNRYNACTGGCCGTCGTTTTACATGGCGTAGCGGGTTCGAGCGCACCGCAGGGTATCCGGCTATTTTTTTTTTTTTTT |
| splint_5pr_lig_idx71 | GTCTCGTGGGCTCGGAGATGTGTATAAGAGACAGCCTGGGAGCATCAGGTAGTAACAGAGTGGTATCNRYNRYNRYNRYNRYNRYNRYNACTGGCCGTCGTTTTACATGGCGTAGCGGGTTCGAGCGCACCGCAGGGTATCCGGCTATTTTTTTTTTTTTTT |
| splint_5pr_lig_idx72 | GTCTCGTGGGCTCGGAGATGTGTATAAGAGACAGTAGCTGACTGTCTTCCATACCGACAGTGGTATCNRYNRYNRYNRYNRYNRYNRYNACTGGCCGTCGTTTTACATGGCGTAGCGGGTTCGAGCGCACCGCAGGGTATCCGGCTATTTTTTTTTTTTTTT |
| splint_5pr_lig_idx73 | GTCTCGTGGGCTCGGAGATGTGTATAAGAGACAGAAGAAACAGGATGACAGAACCCTCAGTGGTATCNRYNRYNRYNRYNRYNRYNRYNACTGGCCGTCGTTTTACATGGCGTAGCGGGTTCGAGCGCACCGCAGGGTATCCGGCTATTTTTTTTTTTTTTT |
| splint_5pr_lig_idx74 | GTCTCGTGGGCTCGGAGATGTGTATAAGAGACAGTACAAGCATCCCAACACTTCCACTAGTGGTATCNRYNRYNRYNRYNRYNRYNRYNACTGGCCGTCGTTTTACATGGCGTAGCGGGTTCGAGCGCACCGCAGGGTATCCGGCTATTTTTTTTTTTTTTT |
| splint_5pr_lig_idx75 | GTCTCGTGGGCTCGGAGATGTGTATAAGAGACAGGACCATTGTGATGAACCCTGTTGTAGTGGTATCNRYNRYNRYNRYNRYNRYNRYNACTGGCCGTCGTTTTACATGGCGTAGCGGGTTCGAGCGCACCGCAGGGTATCCGGCTATTTTTTTTTTTTTTT |
| splint_5pr_lig_idx76 | GTCTCGTGGGCTCGGAGATGTGTATAAGAGACAGATGCTTGTTACATCAACCCTGGACAGTGGTATCNRYNRYNRYNRYNRYNRYNRYNACTGGCCGTCGTTTTACATGGCGTAGCGGGTTCGAGCGCACCGCAGGGTATCCGGCTATTTTTTTTTTTTTTT |
| splint_5pr_lig_idx77 | GTCTCGTGGGCTCGGAGATGTGTATAAGAGACAGCGACCTGTTTCTCAGGGATACAACAGTGGTATCNRYNRYNRYNRYNRYNRYNRYNACTGGCCGTCGTTTTACATGGCGTAGCGGGTTCGAGCGCACCGCAGGGTATCCGGCTATTTTTTTTTTTTTTT |
| splint_5pr_lig_idx78 | GTCTCGTGGGCTCGGAGATGTGTATAAGAGACAGAACAACCGAACCTTTGAATCAGAAAGTGGTATCNRYNRYNRYNRYNRYNRYNRYNACTGGCCGTCGTTTTACATGGCGTAGCGGGTTCGAGCGCACCGCAGGGTATCCGGCTATTTTTTTTTTTTTTT |
| splint_5pr_lig_idx79 | GTCTCGTGGGCTCGGAGATGTGTATAAGAGACAGTCTCGGAGATAGTTCTCACTGCTGAGTGGTATCNRYNRYNRYNRYNRYNRYNRYNACTGGCCGTCGTTTTACATGGCGTAGCGGGTTCGAGCGCACCGCAGGGTATCCGGCTATTTTTTTTTTTTTTT |
| splint_5pr_lig_idx80 | GTCTCGTGGGCTCGGAGATGTGTATAAGAGACAGCGGATGAACATAGGATAGCGATTCAGTGGTATCNRYNRYNRYNRYNRYNRYNRYNACTGGCCGTCGTTTTACATGGCGTAGCGGGTTCGAGCGCACCGCAGGGTATCCGGCTATTTTTTTTTTTTTTT |
| splint_5pr_lig_idx81 | GTCTCGTGGGCTCGGAGATGTGTATAAGAGACAGCCTCATCTTGTGAAGTTGTTTCGGAGTGGTATCNRYNRYNRYNRYNRYNRYNRYNACTGGCCGTCGTTTTACATGGCGTAGCGGGTTCGAGCGCACCGCAGGGTATCCGGCTATTTTTTTTTTTTTTT |
| splint_5pr_lig_idx82 | GTCTCGTGGGCTCGGAGATGTGTATAAGAGACAGACGGTATGTCGAGTTCCAGGACTAAGTGGTATCNRYNRYNRYNRYNRYNRYNRYNACTGGCCGTCGTTTTACATGGCGTAGCGGGTTCGAGCGCACCGCAGGGTATCCGGCTATTTTTTTTTTTTTTT |
| splint_5pr_lig_idx83 | GTCTCGTGGGCTCGGAGATGTGTATAAGAGACAGTGGCTTGATCTAGGTAAGGTCGAAAGTGGTATCNRYNRYNRYNRYNRYNRYNRYNACTGGCCGTCGTTTTACATGGCGTAGCGGGTTCGAGCGCACCGCAGGGTATCCGGCTATTTTTTTTTTTTTTT |
| splint_5pr_lig_idx84 | GTCTCGTGGGCTCGGAGATGTGTATAAGAGACAGGTAGTGGACCTAGAACCTGTGCCAAGTGGTATCNRYNRYNRYNRYNRYNRYNRYNACTGGCCGTCGTTTTACATGGCGTAGCGGGTTCGAGCGCACCGCAGGGTATCCGGCTATTTTTTTTTTTTTTT |
| splint_5pr_lig_idx85 | GTCTCGTGGGCTCGGAGATGTGTATAAGAGACAGAACGGAGGAGTTAGTTGGATGATCAGTGGTATCNRYNRYNRYNRYNRYNRYNRYNACTGGCCGTCGTTTTACATGGCGTAGCGGGTTCGAGCGCACCGCAGGGTATCCGGCTATTTTTTTTTTTTTTT |
| splint_5pr_lig_idx86 | GTCTCGTGGGCTCGGAGATGTGTATAAGAGACAGAGGTGATCCCAACAAGCGTAAGTAAGTGGTATCNRYNRYNRYNRYNRYNRYNRYNACTGGCCGTCGTTTTACATGGCGTAGCGGGTTCGAGCGCACCGCAGGGTATCCGGCTATTTTTTTTTTTTTTT |
| splint_5pr_lig_idx87 | GTCTCGTGGGCTCGGAGATGTGTATAAGAGACAGTACATGCTCCTGTTGTTAGGGAGGAGTGGTATCNRYNRYNRYNRYNRYNRYNRYNACTGGCCGTCGTTTTACATGGCGTAGCGGGTTCGAGCGCACCGCAGGGTATCCGGCTATTTTTTTTTTTTTTT |
| splint_5pr_lig_idx88 | GTCTCGTGGGCTCGGAGATGTGTATAAGAGACAGTCTTCTACTACCGATCCGAAGCAGAGTGGTATCNRYNRYNRYNRYNRYNRYNRYNACTGGCCGTCGTTTTACATGGCGTAGCGGGTTCGAGCGCACCGCAGGGTATCCGGCTATTTTTTTTTTTTTTT |
| splint_5pr_lig_idx89 | GTCTCGTGGGCTCGGAGATGTGTATAAGAGACAGACAGCATCAATGTTTGGCTAGTTGAGTGGTATCNRYNRYNRYNRYNRYNRYNRYNACTGGCCGTCGTTTTACATGGCGTAGCGGGTTCGAGCGCACCGCAGGGTATCCGGCTATTTTTTTTTTTTTTT |
| splint_5pr_lig_idx90 | GTCTCGTGGGCTCGGAGATGTGTATAAGAGACAGGATGTAGAGGGTACGGTTTGAGGCAGTGGTATCNRYNRYNRYNRYNRYNRYNRYNACTGGCCGTCGTTTTACATGGCGTAGCGGGTTCGAGCGCACCGCAGGGTATCCGGCTATTTTTTTTTTTTTTT |
| splint_5pr_lig_idx91 | GTCTCGTGGGCTCGGAGATGTGTATAAGAGACAGGGCTCCATAGGAACTCACGCTACTAGTGGTATCNRYNRYNRYNRYNRYNRYNRYNACTGGCCGTCGTTTTACATGGCGTAGCGGGTTCGAGCGCACCGCAGGGTATCCGGCTATTTTTTTTTTTTTTT |
| splint_5pr_lig_idx92 | GTCTCGTGGGCTCGGAGATGTGTATAAGAGACAGTTGTGAGTGGAAAGATACAGGACCAGTGGTATCNRYNRYNRYNRYNRYNRYNRYNACTGGCCGTCGTTTTACATGGCGTAGCGGGTTCGAGCGCACCGCAGGGTATCCGGCTATTTTTTTTTTTTTTT |
| splint_5pr_lig_idx93 | GTCTCGTGGGCTCGGAGATGTGTATAAGAGACAGAGTTTCCATCACTTCAGACTTGGGAGTGGTATCNRYNRYNRYNRYNRYNRYNRYNACTGGCCGTCGTTTTACATGGCGTAGCGGGTTCGAGCGCACCGCAGGGTATCCGGCTATTTTTTTTTTTTTTT |
| splint_5pr_lig_idx94 | GTCTCGTGGGCTCGGAGATGTGTATAAGAGACAGGATTGTCCTCAAACTGCCACCTACAGTGGTATCNRYNRYNRYNRYNRYNRYNRYNACTGGCCGTCGTTTTACATGGCGTAGCGGGTTCGAGCGCACCGCAGGGTATCCGGCTATTTTTTTTTTTTTTT |
| splint_5pr_lig_idx95 | GTCTCGTGGGCTCGGAGATGTGTATAAGAGACAGCCTGTCTGGAAGAAGAATGGACTTAGTGGTATCNRYNRYNRYNRYNRYNRYNRYNACTGGCCGTCGTTTTACATGGCGTAGCGGGTTCGAGCGCACCGCAGGGTATCCGGCTATTTTTTTTTTTTTTT |
| splint_5pr_lig_idx96 | GTCTCGTGGGCTCGGAGATGTGTATAAGAGACAGCTGAACGGTCATAGAGTCCACCATAGTGGTATCNRYNRYNRYNRYNRYNRYNRYNACTGGCCGTCGTTTTACATGGCGTAGCGGGTTCGAGCGCACCGCAGGGTATCCGGCTATTTTTTTTTTTTTTT |
